# CryoSiam: self-supervised representation learning for automated analysis of cryo-electron tomograms

**DOI:** 10.1101/2025.11.11.687379

**Authors:** Frosina Stojanovska, Ricardo M. Sanchez, Rasmus K. Jensen, Julia Mahamid, Anna Kreshuk, Judith B. Zaugg

## Abstract

Cryo-electron tomography (cryo-ET) enables visualization of macromolecular complexes in their native cellular context, but interpretation remains challenging due to high noise levels, missing information, and lack of ground-truth data. Here, we present CryoSiam (CRYO-electron tomography SIAMese networks), an open-source framework for self-supervised representation learning in cryo-ET. CryoSiam learns hierarchical representations of tomographic data spanning both voxel-level and subtomogram-level information. To train CryoSiam, we generated CryoETSim (CRYO-Electron Tomography SIMulated), a synthetic dataset that systematically models defocus variation, sample thickness, and molecular crowding. CryoSiam trained models transfer directly to experimental data without fine-tuning and support key aspects of cryo-ET data analysis, including tomogram denoising, segmentation of subcellular structures, and macromolecular detection and identification across both prokaryotic and eukaryotic systems. Publicly available pretrained models and the CryoETSim dataset provide a foundation for scalable and automated cryo-ET analysis.

## Introduction

Cryo-electron tomography (cryo-ET) enables three-dimensional (3D) visualization of macromolecules in native cellular environments at nanometer resolution^1,2^. However, interpretation of cryo-ET data is limited by the inherently low signal-to-noise ratio (SNR), missing wedge artifacts, and the heterogeneity of biological specimens^3–5^. Variability in sample thickness, imaging conditions, contrast, and resolution further complicates the analysis, underscoring the need for robust computational methods^5,6^.

Deep learning has emerged as a powerful tool for extracting structural information from cryo-ET data, with the potential to overcome many of the limitations faced by traditional deterministic approaches^7,8^. Yet, most current deep learning approaches rely on supervised learning and require large annotated datasets that are labor-intensive to generate^9–11^. Fully supervised models such as U-Net variants^12^ (e.g., DeepFinder^13^ and DeePiCt^14^) achieve strong segmentation performance, but require voxel-level annotation, and often fail to generalize across tomograms with varying contrast and SNR^13–15^. Recent advances in semi-supervised and self-supervised learning (e.g., TomoTwin^16^, MiLoPYP^17^, and CryoSAM^18^) have improved generalization while reducing dependence on manual annotation, illustrating the potential of representation learning for macromolecular localization and identification in tomograms^16–19^. However, these methods still rely on experimental training data or task-specific supervision, and often fail to transfer effectively across datasets with differing imaging conditions or structural compositions without fine tuning.

A persistent challenge in developing deep learning methods for cryo-ET is the lack of complete and reliable ground truth for model training and evaluation^14,16^. While recent community efforts have produced curated experimental datasets with voxel-level annotations^20^, such data remain limited in scope and molecular diversity. As experimental tomograms lack comprehensive voxel-level labels, simulated data offer a controlled alternative^21–24^. However, existing simulation datasets, such as SHREC^23^, fall short of representing the molecular crowdedness and structural heterogeneity, contrast and defocus variations that are characteristic of real cellular tomograms. These discrepancies limit their utility as ground-truth data or benchmarks^22,25^, and underscores the need for a more representative, large-scale simulated dataset that better captures the diversity of experimental cryo-ET data^21,26^.

To address this limitation, we developed CryoETSim (CRYO-Electron Tomography SIMulated dataset), a synthetic dataset generated using the cryo-TomoSim (CTS) simulator^21^. CryoETSim models tomograms at consistently high molecular crowding while systematically varying parameters such as sample thickness and defocus, reproducing the modulation and contrast properties observed in experimental data. Our dataset comprises 400 tomograms covering a broad range of molecular structures and imaging conditions, providing sufficient scale and diversity to train robust deep learning models for cryo-ET analysis.

Building on this foundation, we introduce CryoSiam (CRYO-electron tomography SIAMese networks), an open-source framework trained in a self-supervised manner using the SimSiam embedding paradigm^27,28^, designed to learn representations at both the voxel and the subtomogram level. Unlike standard natural-image transformations required for self-supervised learning, which assume scale or color invariance^29^, CryoSiam employs transformations tailored to the physical and statistical characteristics of cryo-ET, including defocus variation, frequency filtering, and noise modeling.

We show that CryoSiam, trained exclusively on the CryoETSim dataset, demonstrates cross-domain generalization when applied directly to experimental tomograms. We evaluated its performance in denoising, segmenting, and identifying macromolecular structures using publicly available cryo-ET data from a minimal bacterium (*Mycoplasma pneumoniae*) and a eukaryotic single-cell alga (*Chlamydomonas reinhardtii*), covering a broad range of biological complexity, sample properties, and imaging conditions. By providing pretrained models, CryoSiam establishes a practical foundation for automated, scalable cryo-ET analysis and systematic exploration of macromolecular organization *in situ*.

## Results

### Overview of the CryoSiam framework

CryoSiam is a deep learning framework designed to extract structural information from in-cell cryo-ET data (Fig. 1). The framework consists of two complementary self-supervised modules that operate at different spatial scales: DenseSimSiam (DENSE SIMple SIAMese networks) at the voxel level and SimSiam (SIMple SIAMese networks) at the subtomogram level. Together, they learn multi-scale embedding representations that capture both local and contextual structural features.

**Figure 1:**
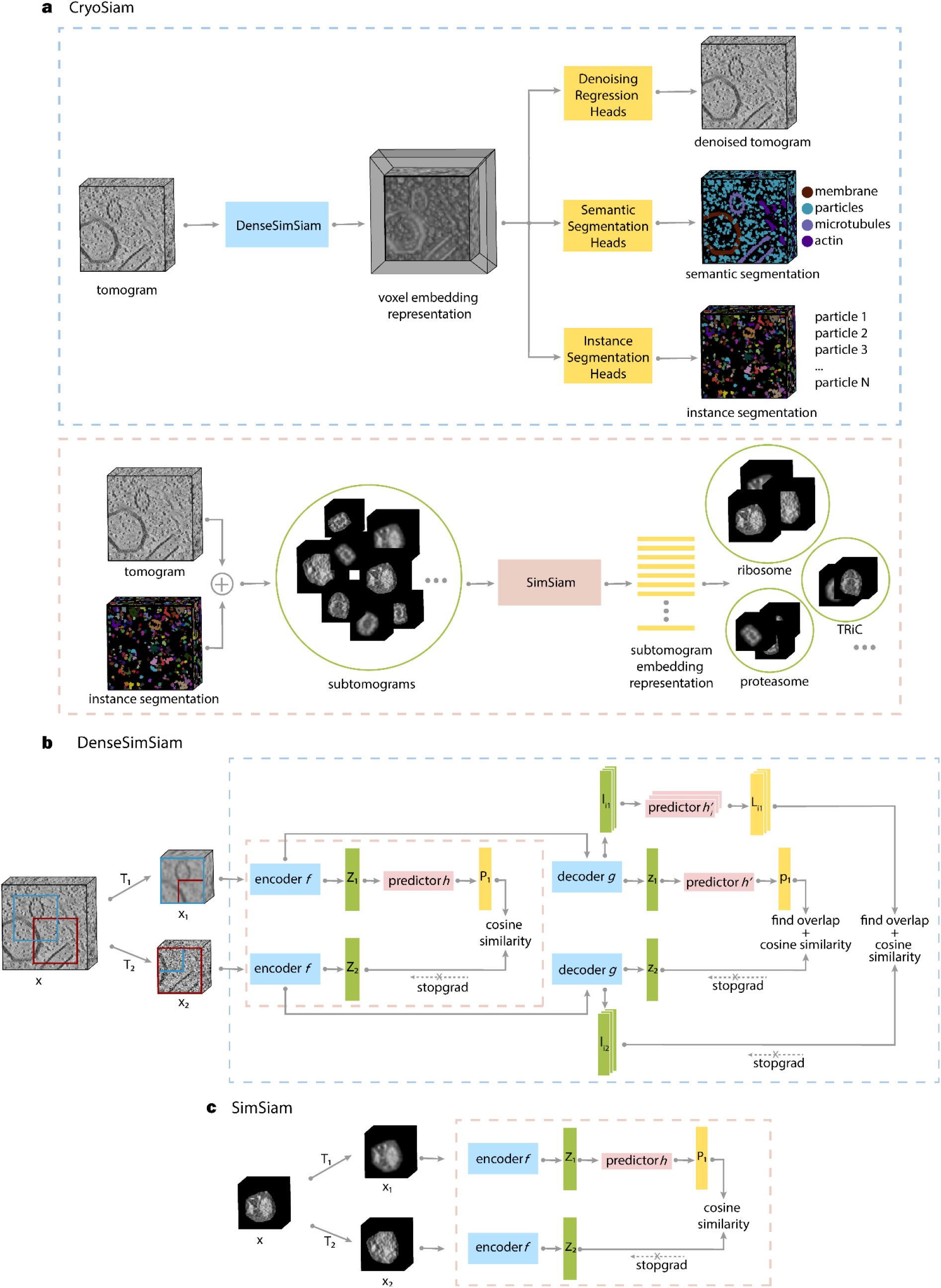
Overview of the CryoSiam framework for self-supervised representation learning of in-cell cryo-ET data. **a**. The CryoSiam pipeline integrates two complementary self-supervised modules for multi-scale representation learning. DenseSimSiam (top) processes input tomograms to generate voxel-level embeddings used by task-specific heads for tomogram denoising, semantic segmentation (membrane, particle, microtubule, actin, and DNA/RNA filament classes), and instance segmentation (individual particle masks). Subtomograms extracted using the instance segmentation masks are then processed by SimSiam (bottom), which produces subtomogram-level embeddings capturing macromolecular features for downstream analyses such as macromolecular identification. **b**. Architecture of the DenseSimSiam module. Overlapping augmented patches are sampled from tomograms and passed through a shared encoder *f* to generate embeddings of the full patch. A decoder *g* based on a Feature Pyramid Network (FPN) reconstructs hierarchical features, while predictors (*h*′ and *h*′_*i*_) combined with stop-gradient operations prevent representational collapse. Training optimizes cosine similarity between embeddings from overlapping patches at the patch, voxel, and FPN levels. **c**. Architecture of the SimSiam module. Subtomograms obtained from instance masks undergo two distinct transformations (*T*_1_ and *T*_2_) and are processed by an encoder *f* and predictor *h* with a stop-gradient operation. Cosine similarity between embeddings from the two views enforces self-supervised learning of stable, discriminative subtomogram representations.

DenseSimSiam processes 3D tomograms to produce voxel-wise embeddings (Fig. 1a). These embeddings form a dense feature space that enables various voxel-wise downstream tasks via task-specific heads. We implemented tomogram denoising, semantic segmentation, and instance segmentation within the CryoSiam framework. For the denoising task, the model predicts a noise-reduced tomogram to enable direct visualization of structural details. Semantic segmentation assigns each voxel to one of several structural classes: membrane, particles (i.e. macromolecular complexes), microtubules, actin, thin filaments (e.g., DNA and RNA), or background. The instance segmentation isolates and distinguishes individual particles. These segmented particle instances are further embedded into representations that reflect their structural similarity. For this, instance masks from the segmentation stage are used to guide subtomogram extraction, defining the spatial boundaries of each particle and masking surrounding density. The resulting cropped subtomograms are then processed by the SimSiam module (Fig. 1a), which learns embeddings representing the structural and local contextual characteristics of individual particles through self-supervised learning. To further improve discrimination between distinct structural representations, contrastive learning was applied in a subsequent stage.

The self-supervised learning strategy in the DenseSimSiam module builds on Siamese network-based methods for representation learning^27,28^. Briefly, two partially overlapping 3D patches, *x*_1_ and *x*_2_, are randomly sampled from a tomogram *x*. The model is trained to enforce similarity between voxel embeddings from the overlapping regions of both patches (Fig. 1b), promoting spatial and contextual consistency in the learned representations. Distinct transformations *T*_1_ and *T*_2_, specifically chosen to capture variation in cryoET data (Extended Data Fig. 1, Methods), are applied to each patch before they are passed through a shared encoder *f*, composed of a ResNet^30^ backbone followed by a multi-layer perceptron (MLP) projection head (Fig. 1b). The encoder outputs two embeddings, *Z*_1_ and *Z*_2_, corresponding to the transformed patches *x*_1_ and *x*_2_.

To prevent representation collapse during training, we include a fully connected predictor *h* and apply a stop-gradient operation as described previously^27^. The predictor generates a full-patch embedding *P*_1_, while a decoder *g*, based on a Feature Pyramid Network (FPN)^31^, reconstructs voxel-level embeddings *z*_1_ and *z*_2_, and produces intermediate embeddings from each level *i* in the FPN, denoted *I*_*i*1_ and *I*_*i*2_. Similar to the full-patch embeddings, predictors *h*′ and *h*′_*i*_, along with stop-gradient operations, are used during training to prevent collapse. Finally, a joined cosine similarity loss is computed between the embeddings from overlapping regions at patch, voxel and FPN levels, encouraging the model to learn consistent representations across spatial resolutions (Fig. 1b).

While DenseSimSiam learns voxel-level features that capture local structural context, the complementary SimSiam module operates at the subtomogram level to model higher-order representations of individual macromolecular complexes. In this module, two distinct, cryo-ET-specific, transformations, *T*_1_ and *T*_2_ (Methods), are applied to a subtomogram *x* to generate augmented views *x*_1_ and *x*_2_ (Fig. 1c). The model is trained to produce similar embeddings for both transformed inputs, encouraging the network to learn invariant and stable representations of macromolecular structures. The encoder *f*, composed of a ResNet backbone followed by a MLP projection head, outputs embeddings that are optimized using a cosine similarity loss function, aligning representations from the two transformed views. To prevent training collapse, a predictor *h* and a stop-gradient operation are incorporated^27^.

Through this training strategy, the SimSiam module learns compact and discriminative subtomogram embeddings that capture the structural characteristics of individual macromolecular complexes while remaining robust to variations in orientation, noise, and contrast. These representations provide a higher-level description of particle structure, complementing the voxel-level features learned by DenseSimSiam, and form the basis for downstream analyses such as macromolecular identification.

### Generation of a cryo-ET simulation dataset with realistic characteristics

Ground truth data are essential for developing and evaluating deep learning methods, but obtaining complete voxel-level annotations from experimental cryo-ET data is impossible. Although several annotated experimental datasets exist, they typically cover only a small subset of well-characterized structures and lack comprehensive voxel-level labeling^14^. Available simulated datasets^23^ partially address this limitation, but remain limited with regard to diversity of molecular species and fail to reproduce the molecular crowdedness, contrast variation, and structural complexity of real tomograms. As a result, models trained on such data struggle to transfer effectively to real tomograms. Recent advances in tomogram simulation have substantially improved the ability to reproduce realistic imaging conditions and molecular crowdedness, with the CTS simulator^21^ providing a particularly flexible framework for generating high-fidelity synthetic tomograms.

Building on this progress, we generated the CryoETSim dataset (Supplementary Note 1, Methods). CryoETSim comprises 400 tomograms, representing three biologically distinct sample types (Supplementary Table 1). The general sample includes 141 macromolecular structures similar to those used in previous studies^16,21^ spanning molecular weights from 10 kDa to 4000 kDa, together with structural elements such as membranous vesicles, membrane-embedded complexes, microtubules and actin filaments (Fig. 2a, Supplementary Table 2). To emulate realistic molecular crowding, particles were sequentially added from largest to smallest, increasing particle number as molecular size decreased and reaching crowding densities of up to 80% (Fig. 2b).

**Figure 2:**
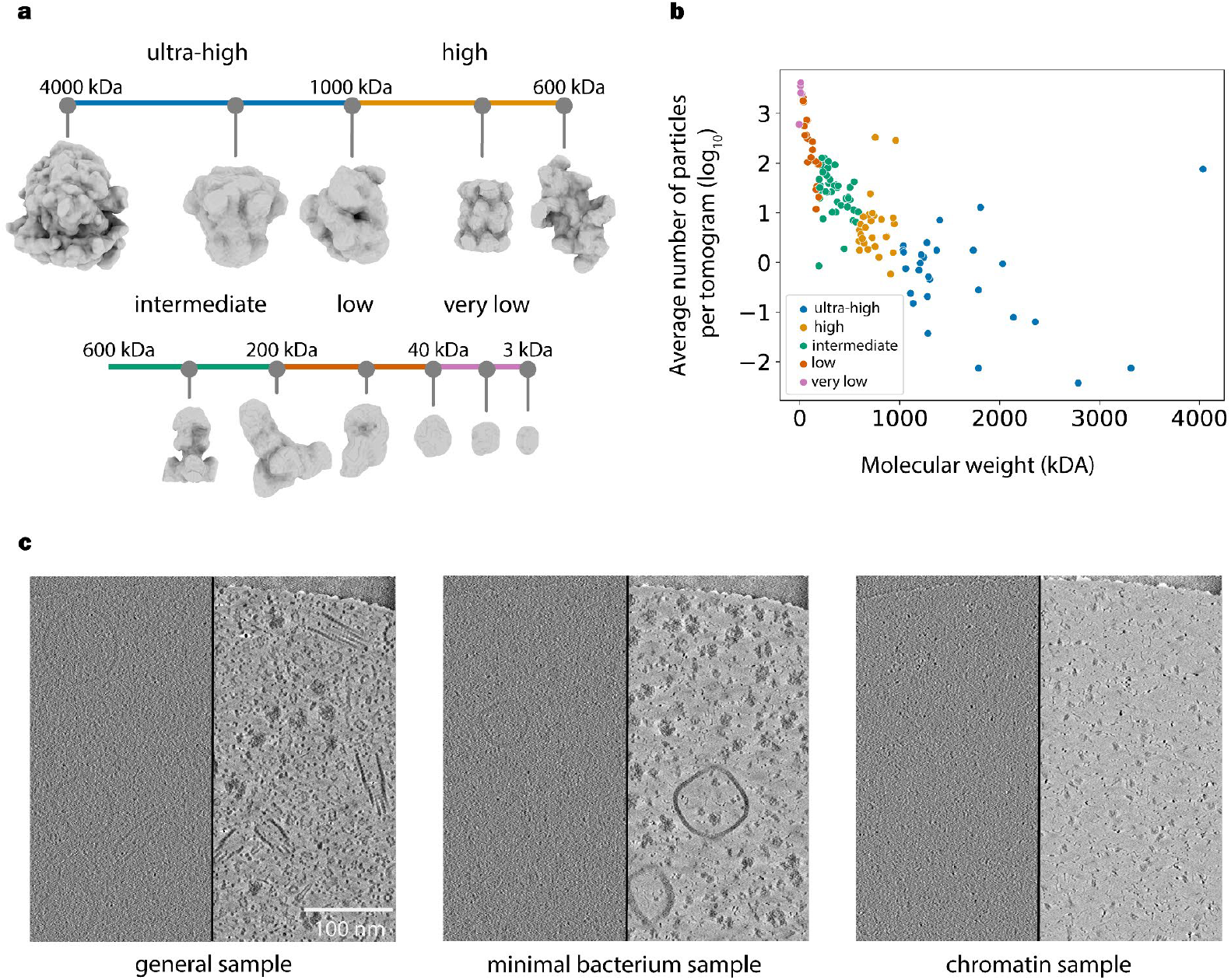
Simulation of tomograms with varying particle sizes, defocus, and sample thickness. **a**. Classification of particles by molecular weight into six size categories: ultra-high (>1000 kDa), high (1000-600 kDa), intermediate (600-200 kDa), low (200-40 kDa), and very low (<40 kDa) molecular weight. Representative particle maps from each category are shown at the same scale to illustrate their size and variable morphology. **b**. Scatter plot showing the relationship between molecular weight and the average particle number per tomogram (log scale) for each size category. Data points are color-coded by particle size categories (**a**), revealing trends in particle abundance across molecular weight classes. **c**. Representative simulated tomograms for three sample types: general, minimal bacterium, and chromatin. Each panel shows an unmodulated simulation on the right and a noisy simulated tomogram with CTF modulation on the left, highlighting the effect of noise and optical modulation on contrast and particle visibility.

Two additional specialized datasets were generated: (1) a minimal bacterium sample, representing mainly the transcription and translation machineries with complete ribosomes, large and small ribosomal subunits, RNA polymerase, smaller cytosolic complexes, and simulated DNA filaments (Supplementary Table 2); and (2) a chromatin sample, composed of nucleosomes and DNA filaments ranging from 50 to 1000 base pairs, modeled using a wormlike-chain (WLC) approach^32^ to reproduce realistic folding and flexibility (Fig. 2c, Supplementary Fig. 1). These specialized datasets provide complementary biological contexts: the minimal bacterium sample represents a system with well-defined macromolecular assemblies essential for gene expression, while the chromatin sample mimics a less crowded environment resembling isolated biomolecular assemblies.

Tomograms were simulated using the CTS software^21^. Sample thicknesses ranged from 35 nm to 100 nm, where thinner samples displayed higher contrast (Extended Data Fig. 2a). Modeling thicker specimens (>100 nm) was outside the scope of this study, but will be explored in future extensions of the dataset. Each 3D model simulated on 6.8 Å/pixel was projected into a synthetic tilt series, modulated by a contrast transfer function (CTF) with varying defocus values, corrupted with smoothing and additive Gaussian noise, and reconstructed back to 3D (Methods). To reproduce the imaging variability of experimental tomograms, defocus values ranged from 2 µm, which enhances high-frequency information and yields sharper images, to 5 µm, which emphasizes low-frequency components and increases contrast (Extended Data Fig. 2b). For each simulation, we generated both the clean tomogram (reconstructed from unmodulated projections) and the noisy, CTF-modulated tomogram, allowing direct comparison between ideal and realistic experimental conditions. Together, these parameters capture the visual modulation, contrast variation, and structural diversity characteristic of experimental cryo-ET data, providing a realistic foundation for training and evaluating CryoSiam.

### Voxel-level structural representation learning with DenseSimSiam

The DenseSimSiam model was trained in a self-supervised manner on the clean tomograms from the CryoETSim dataset. To promote robust representation learning and enhance feature invariance to common distortions in cryo-ET data, we carefully designed the two transformations *T*_1_ and *T*_2_ (Fig.1b). These transformations included random Gaussian low-pass and high-pass filtering, which simulate frequency-dependent signal attenuation, additive Gaussian noise, and random voxel masking, which encourages the network to infer missing structural context (Extended Data Fig. 1). Together, these transformations enable the model to learn representations that are resilient to variations in contrast, noise, and molecular density. We further optimized the network architecture and training parameters through systematic ablation studies to obtain the final CryoSiam configuration (Supplementary Table 6-11).

Following training, DenseSimSiam generates 64-dimensional voxel-level embedding representations for an input tomogram, encoding both local structural context and spatial relationships. To visualize this high-dimensional embedding space, we applied principal component analysis (PCA) to reduce the embeddings to three dimensions and mapped the resulting components to RGB color channels (Fig. 3a). In this representation, voxels with similar learned features appear in similar colors, and the resulting spatial pattern closely follows the structures visible in the clean tomogram, including membranes, microtubules and macromolecular complexes. This demonstrates that DenseSimSiam captures meaningful structural information directly from tomographic data without supervision.

**Figure 3:**
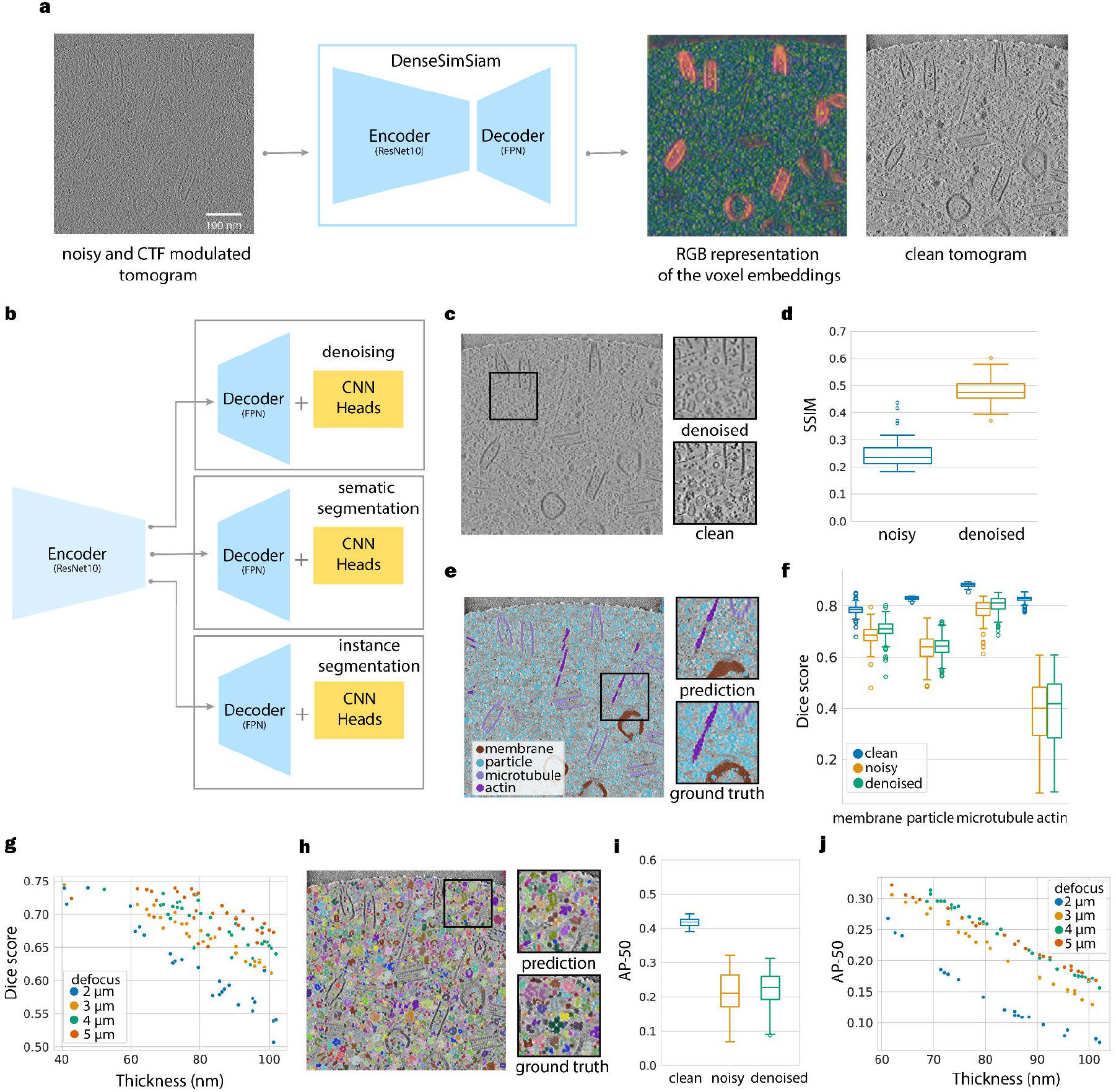
Evaluation of DenseSimSiam performance with simulated data on various downstream tasks. **a**. Voxel embedding generation. Noisy tomograms were processed with the trained DenseSimSiam model to generate voxel embeddings. Principal component analysis (PCA) was applied to reduce embedding dimensionality, with the first three components mapped to RGB channels. The resulting color representation highlights distinct structural features and aligns with the corresponding clean tomogram. **b**. Training of downstream tasks. DenseSimSiam serves as a self-supervised backbone for multiple tasks. The encoder was frozen, and the decoder, equipped with task-specific CNN heads, was trained independently for tomogram denoising, semantic segmentation, and instance segmentation. **c**. Example of tomogram denoising. The same noisy tomogram slice shown in panel a) is compared to its denoised and clean counterparts. **d**. Structural similarity index (SSIM) for denoising, calculated relative to the clean tomograms. **e**. Qualitative results for semantic segmentation. Predictions from the model are compared to the ground truth for various structural components, including membranes, particles, microtubules, and actin filaments. **f**. Dice scores, computed against the clean ground truth masks, assess performance across structural components, including membranes, particles, microtubules, and actin filaments. **g**. Impact of sample thickness and defocus on semantic segmentation. Dice scores are computed relative to the clean ground truth. **h**. Instance segmentation performance. Model predictions are compared to the instance-level ground truth masks. **i**. AP-50 evaluation for instance segmentation across clean, noisy, and denoised tomograms. **j**. Influence of sample thickness and defocus on instance segmentation.

Once trained, DenseSimSiam serves as a self-supervised backbone for a range of downstream tasks. For this, the encoder-decoder architecture is extended with task-specific convolutional neural network (CNN) heads, each consisting of two convolutional layers (Fig. 3b). During the training of the downstream tasks, the ResNet10 encoder remains frozen to preserve the learned representations, while the weights of the feature pyramid network (FPN) decoder and task-specific heads are optimized for each task. This strategy enables efficient fine-tuning for tomogram denoising, semantic segmentation, and instance segmentation while maintaining the robustness of the pretrained backbone. The model was trained and evaluated on the CryoETSim dataset. We used 280 tomograms (from all 3 simulated sample types) for training the denoising and segmentation tasks and 200 tomograms for the instance segmentation (10% used for validation during training), while reserving 119 and 100 tomograms for testing the two tasks, respectively (Supplementary Table 3). Because full tomograms exceed GPU memory capacity, both training and inference were performed on 3D patches of 128^3^ voxels, extracted using a sliding-window approach and subsequently reassembled into complete tomograms.

The tomogram denoising task was evaluated using noisy, CTF-modulated tomograms as inputs and the corresponding clean tomograms as targets. The denoised tomogram, derived from the noisy tomogram (Fig. 3a), exhibited improved structural definition, recovering fine features such as membranes, filamentous structures and particles (Fig. 3c). Quantitatively, performance was assessed using the structural similarity index metric (SSIM), which measures local contrast and structural correspondence between the denoised and clean tomograms. A SSIM score of 1 means perfect similarity, 0 no similarity and −1 reflects negative similarity. SSIM scores increased consistently after denoising, confirming that the model effectively reduces noise while maintaining overall structural fidelity (Fig. 3d). However, the values did not approach unity because the simulated noise is inherently mixed with the signal prior to CTF modulation and tomogram reconstruction, creating a complex interaction that cannot be fully disentangled. As a result, the denoising task must balance suppressing noise with preserving genuine structural features, leading to slightly smoother reconstructions compared with the clean tomograms (Fig. 3c).

The semantic segmentation task involved classifying five biologically distinct classes: membranes, particles, microtubules, actin, and background (Fig. 3e). For the minimal bacterium and chromatin samples, an additional filament class was included to represent short DNA and RNA filaments. To evaluate the influence of noise and CTF modulation on the performance of semantic segmentation, three models were trained separately on (i) clean tomograms, (ii) noisy, CTF-modulated tomograms, and (iii) denoised tomograms obtained as predictions from the preceding denoising task. Overall, the models achieved high segmentation accuracy on clean data, with mean Dice scores above 0.7 for all of the classes (Fig. 3f; Supplementary Table 4). Performance declined on noisy tomograms, reflecting the expected impact of reduced SNR, but the degree of robustness varied across structures. Microtubules showed the highest resilience to noise and contrast variation, followed by membranes and particles, whereas actin exhibited a pronounced performance drop, highlighting the particular sensitivity of thin filamentous features to noise and modulation artifacts (Fig. 3f; Supplementary Table 4). The DNA/RNA filament class, which was less represented in the simulated tomograms, had a similar Dice score to actin and was also strongly affected by noise (Supplementary Table 4). Denoising improved segmentation performance only marginally compared to the noisy tomograms, indicating that noise removal enhances local contrast but does not fully restore the lost information.

We next examined how defocus and sample thickness jointly affect segmentation performance on the noisy tomograms. As expected, we found that Dice scores consistently decreased with increasing sample thickness, reflecting reduced SNR in thicker samples. Similarly, lower defocus values yielded weaker contrast at a given thickness, further diminishing segmentation accuracy (Fig. 3g).

For structural analysis of macromolecular complexes in crowded tomograms, accurate separation of individual particles is essential. CryoSiam uniquely enables this by performing instance-level segmentation, predicting voxel-wise masks that assign each voxel to a distinct object rather than a general class. This capability bridges voxel-level representation learning with downstream structural tasks such as subtomogram embedding. To achieve this, we trained instance segmentation heads to predict three voxel-wise maps: foreground probability, distance to background, and instance boundary probability (Supplementary Fig. 2a). Particle markers were extracted from the distance-to-background map after Gaussian smoothing and adaptive thresholding. A seeded watershed algorithm^33^ was then applied on the distance-to-background map, masked by the foreground prediction, to generate preliminary instance segmentation labels. To correct for over-segmentation, a multicut refinement^34,35^ was performed using the watershed instances together with the predicted boundary map, merging erroneously split regions and improving particle separation in the crowded simulations (Supplementary Fig. 2a).

The resulting masks successfully separated neighboring particles even in densely packed regions, with remaining errors primarily occurring in low-contrast areas and within hollow structures (Fig. 3h). Performance was evaluated using average precision (AP) across intersection-over-union (IoU) thresholds, with AP-50 (IoU = 0.5) used as the main metric (Fig. 3i; Supplementary Table 5). When both training and prediction were performed on denoised tomograms, models achieved slightly higher AP-50 scores compared to those trained and predicted on noisy data, indicating a modest benefit from denoising. Segmentation accuracy decreased with increasing sample thickness and lower defocus values, consistent with reduced contrast and structural separability under these imaging conditions (Fig. 3j). Furthermore, larger particles achieved higher AP scores across all conditions, though the relationship with size was not strictly monotonic, with the greatest gains of prior denoising observed for large particles (Supplementary Fig. 2b).

In conclusion, across all downstream tasks, DenseSimSiam demonstrated that representations learned through self-supervised training on simulated data effectively generalize across the simulated dataset. The model enhanced tomogram interpretability through denoising, segmented major structural components, and reliably separated individual macromolecules even in densely crowded regions. These results show that DenseSimSiam captures meaningful voxel-level features that are adaptive to imaging variability and noise, providing a strong foundation for downstream analysis of cryo-ET data. In practical terms, DenseSimSiam provides the location and a mask for each particle, which can be used for further analysis.

### Subtomogram-level representation learning with SimSiam

The second component of the CryoSiam framework, SimSiam, learns subtomogram-level representations that are designed to capture the structural identity of individual particles. For training, we again used the CryoETSim dataset and extracted up to ten subtomograms per structural class per tomogram to mitigate dataset imbalance caused by the higher abundance of small particles. Because the molecular crowdedness of tomograms introduces strong contextual interference, subtomograms were first extracted using the ground-truth instance masks to separate individual particles. Each subtomogram was cropped around the corresponding particle based on its instance mask and then zero-padded to a uniform volume of 64^3^ voxels to ensure consistent input dimensions. A loose convex-hull mask derived from each instance was then applied to remove surrounding densities and background voxels while preserving the overall particle shape. Training used 200 tomograms from the general CryoETSim sample, with 10% reserved for validation, and 100 tomograms for testing, yielding 175,371 and 88,585 subtomograms, respectively.

SimSiam was trained in a self-supervised manner using the same Gaussian low-pass, high-pass, and noise augmentations defined for DenseSimSiam, together with affine and cropping transformations that introduced random rotations, translations, and partial cropping. A UMAP^36^ projection of the 1024-dimensional learned embeddings, colored by the log_10_ of the instance mask volume, showed that particles of different molecular sizes occupy distinct regions of the representation space (Fig. 4a). Classification based on these embeddings using a Multi-Layer Perceptrons (MLP) or k-nearest neighbors (k-NN) classifier achieved median F1 scores above 0.94 on clean data, with moderate reductions under noise and partial recovery after denoising (Fig. 4b). The MLP classifier slightly outperformed k-NN, suggesting that non-linear feature relationships, only captured by MLP, improve prediction.

**Figure 4:**
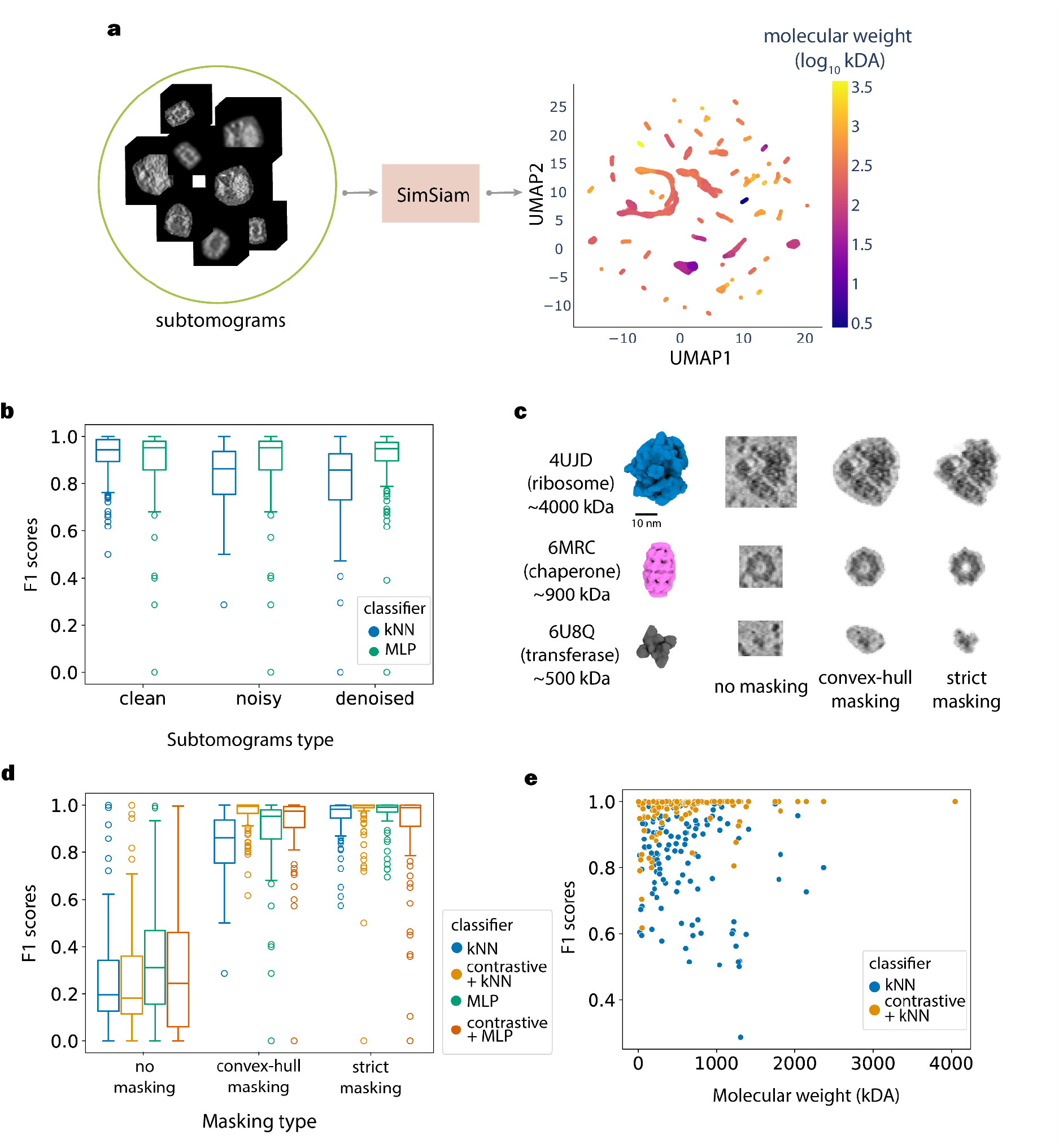
Evaluation of SimSiam-based subtomogram representation learning and classification performance. **a**. Subtomograms were cropped around particles using ground truth instance masks from the CryoETSim dataset and masked with loose convex-hull masks to remove surrounding densities. Each subtomogram was processed by the self-supervised SimSiam model to generate embeddings, which were projected into 2D using UMAP and color-coded by molecular weight (log_10_ scale, kDa). **b**. Classification performance across data conditions. Subtomogram embeddings from clean, noisy, and denoised data were classified using k-nearest neighbors (kNN) and a multilayer perceptron (MLP). **c**. Masking strategies for subtomogram extraction. Representative subtomograms of three particle types (ribosome, chaperone, and transferase; corresponding PDBs provided) are shown without masking, and with convex-hull or strict masking. **d**. F1 score distributions for kNN and MLP classifiers under the three masking strategies, with additional comparisons between standard SimSiam embeddings and those refined with contrastive learning. **e**. Molecular weight dependence under convex-hull masking. F1 scores plotted as a function of molecular weight for noisy subtomograms processed with convex-hull masking, and with contrastive refinement.

To test whether molecular crowding and background noise affect the ability of SimSiam to learn discriminative representations, we compared three masking strategies: no masking, strict masking, and convex-hull masking (Fig. 4c). In the no-masking approach, the subtomogram is cropped with dimensions from the bounding box that includes the particle (padded with zeros to have a 64^3^ cube), retaining background noise and neighboring structures. This obscures relevant features, particularly for smaller particles, even when the cropped volume was adjusted to the particle size, rather than cropped by a fixed-size cube (Fig. 4d). Strict masking removed all voxels outside the ground-truth instance mask, isolating particles completely but occasionally truncating peripheral densities of complex or elongated structures. To balance these effects, convex-hull masking was introduced as a loose geometric enclosure around each particle, reducing background interference while preserving overall shape. Classification performance on noisy subtomograms showed a strong dependence on masking strategy (Fig. 4d). Unmasked subtomograms yielded the lowest F1 scores, whereas convex-hull and strict masking substantially improved performance, resulting in a clear increase in discriminative power. Convex-hull masking provided the optimal balance between removing neighbourhood context and allowing imperfect predicted particle masks. These results further highlight the importance of accurate instance segmentation, such as that provided by the DenseSimSiam part of CryoSiam, for effective representation learning in densely-populated tomograms.

To further refine the learned representations, we applied supervised contrastive learning as an additional training stage and reassessed performance using the k-NN classifier, which determines particle identity based on cosine similarity to embeddings in the training set. Unlike the MLP classifier, which learns a more complex decision function, k-NN directly reflects the structure of the embedding space, providing an interpretable measure of representational quality. Comparing embeddings from the initial SimSiam model with those refined through contrastive learning on the noisy subtomograms revealed consistent F1 score improvements across masking strategies (Fig. 4d), demonstrating that contrastive fine-tuning enhances representation separability and robustness to noise. Notably, classification performance remained uniformly high across molecular weights, with F1 scores ranging from approximately 0.85 to above 0.95 for most particle classes. The relationship between particle size and performance was not strictly linear, as several small complexes also achieved F1 scores above 0.95 (Fig. 4e). The confusion matrix further confirms consistent separation between molecular classes and limited cross-class misassignment (Supplementary Fig. 3). This highlights that, if these particles can be reliably isolated as individual instances from the noisy tomograms, they can be classified equally well as large particles. Overall, these results demonstrate that contrastive learning strengthens the discrimination of macromolecular identities within subtomogram embeddings and further underscores the importance of accurate instance-based masking strategies for representation learning in crowded tomograms.

Together, these results demonstrate that SimSiam learns robust, discriminative representations of macromolecular structures at the subtomogram level, capable of separating molecular identities even under high crowding and noise and low molecular weight, forming a foundation for reliable transfer to experimental data.

### Voxel-level analysis of experimental cryo-ET data

To evaluate the generalization of CryoSiam to experimental data, models trained exclusively on simulated tomograms were directly applied to cryo-electron tomograms of *Mycoplasma pneumoniae*^*37*,38^ (EMPIAR-10499) and *Chlamydomonas reinhardtii*^*39*^ (EMPIAR-17756) cells. These two datasets represent respectively prokaryotic and eukaryotic cell types, span different microscope and detector generations, imaging conditions, and sample preparation approaches (Supplementary Table 12). The CryoSiam framework was applied to these datasets for tomogram denoising, semantic segmentation, and instance segmentation.

CryoSiam effectively denoised the experimental tomograms, enhancing structural visibility while preserving native biological features (Fig. 5a). Compared to Gaussian filtering, which oversmooths and suppresses fine details, CryoSiam restored sharp structural boundaries and improved overall contrast while reducing high-frequency noise. A comparison with the Noise2Noise-based Cryo-CARE^40^ method on *C. reinhardtii* tomograms (Extended Data Fig. 3a) showed that CryoSiam also improved contrast that was lost due to modulation by the CTF, reflecting its training on clean simulated data. Despite being trained solely on simulated data, CryoSiam generalized effectively to experimental tomograms, capturing realistic ultrastructural features and demonstrating strong transfer performance across biological systems.

**Figure 5:**
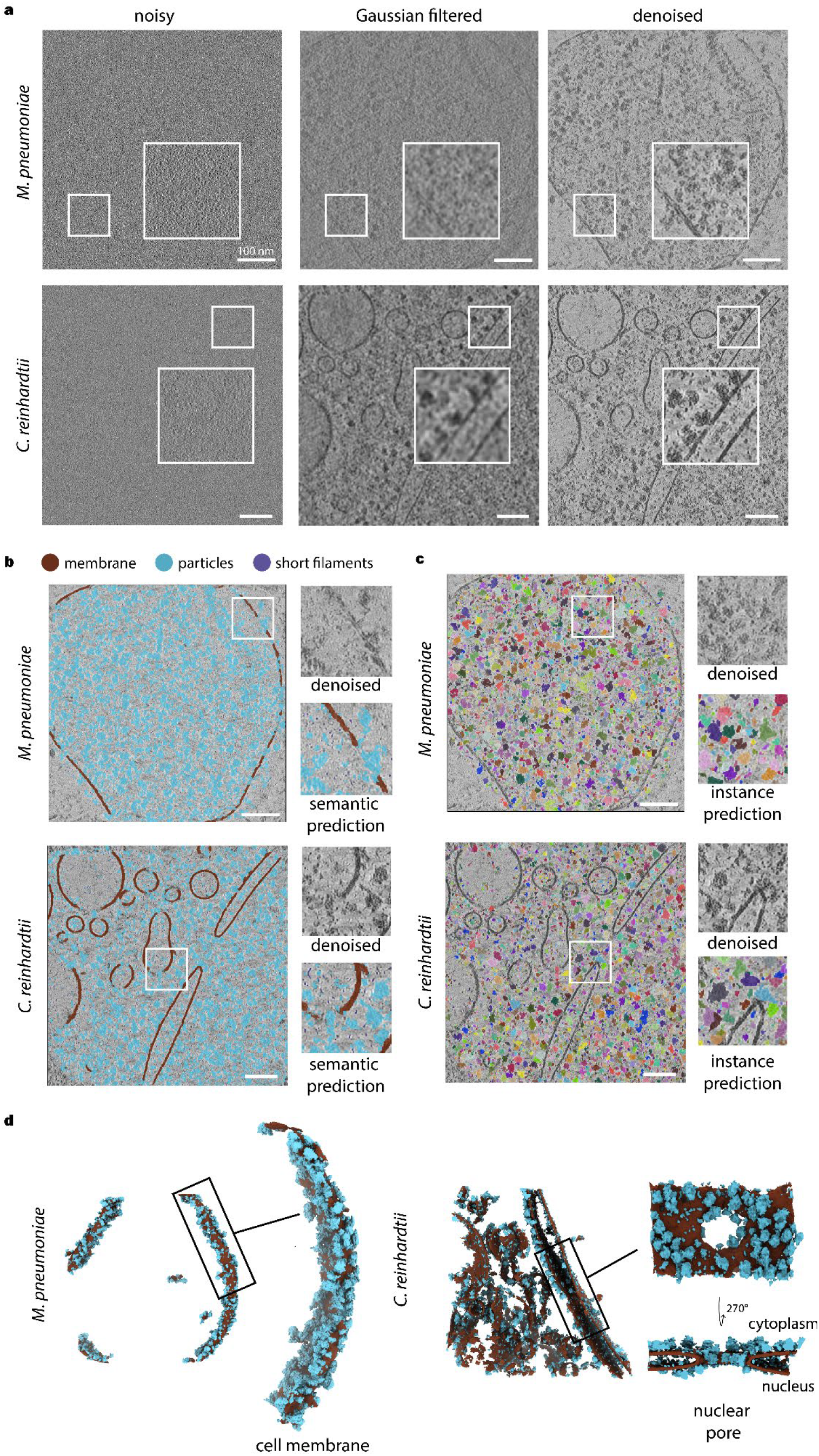
Application of CryoSiam to experimental cryo-electron tomograms of *Mycoplasma pneumoniae* and *Chlamydomonas reinhardtii*. **a**. Tomogram denoising. CryoSiam models trained on simulated data were directly applied to experimental tomograms from two distinct organisms, the prokaryote *M. pneumoniae* and the eukaryote *C. reinhardtii*. **b**. Semantic segmentation. Using the denoised tomograms together with the predicted cell/lamella mask, CryoSiam accurately identifies major cellular components, including membranes (brown), particles (blue), and thin filaments (purple). **c**. Instance segmentation. Particle instance predictions reveal individual macromolecular complexes throughout the samples. Insets show comparisons between denoised tomograms and instance masks. **d**. Integrated membrane-particle visualization. Semantic membrane predictions were combined with instance masks of particles located in contact with or within one to two voxels of the membrane surface, enabling visualization of macromolecular organization at membrane interfaces. In *C. reinhardtii*, the nuclear pore was visualized with an additional sliced view in the middle.

An additional downstream task predicted 3D cell/lamella masks to distinguish the region of the physical sample within the overall reconstructed tomogram (Extended Data Fig. 3b). Trained on denoised simulated tomograms and applied to denoised experimental data, this prediction effectively excluded regions containing only noise or reconstruction artifacts, improving the precision of subsequent analyses. Using the denoised tomograms and lamella masks as input, CryoSiam (semantic segmentation mode) accurately delineated membranes (brown) and particles (blue) in experimental data (Fig. 5b). Although trained only on short simulated DNA/RNA filaments, the network also identified filamentous densities (purple) in real tomograms, indicating that CryoSiam captures generalizable structural features rather than dataset-specific patterns.

Subsequent instance segmentation further refined the analysis to the level of individual macromolecular complexes (Fig. 5c). CryoSiam effectively separated distinct particles within crowded intracellular regions. Insets highlight detailed instance-level predictions overlaid on the denoised tomograms, confirming that the model can differentiate closely spaced macromolecules without fine-tuning on experimental data. This ability to generate accurate instance masks is critical for downstream analyses, providing the structural delineation required for subtomogram embedding and exploration with the SimSiam component.

To illustrate the combined interpretive power of these multilevel predictions, membrane segmentations were integrated with nearby particle instances within two voxels of the membrane surface (Fig. 5d). This integrated visualization revealed the organization of macromolecules at membrane interfaces. When compared with predictions from the supervised membrane segmentation method MemBrain-Seg^41^ (Extended Data Fig. 3c), CryoSiam achieves comparable membrane delineation quality while additionally providing spatial information on nearby macromolecules through its instance segmentation task. In *M. pneumoniae*, the reconstructed cell membrane prediction is shown together with the associated particle instances, revealing densely arranged macromolecules embedded in or adjacent to the membrane. A magnified view highlights the local architecture of membrane-associated complexes and their spatial distribution relative to the membrane surface (Fig. 5d). In *C. reinhardtii*, a similar analysis of the nuclear envelope membrane shows the arrangement of predicted macromolecular complexes along the membrane and within the nuclear pore (Fig. 5d).

### Subtomogram embeddings and molecular identification in experimental tomograms

To explore whether CryoSiam representations capture biologically meaningful features in experimental data, we analyzed the instance segmentation predictions from *M. pneumoniae* and *C. reinhardtii* tomograms. Predicted particle instances were cropped using convex-hull masks to isolate individual macromolecules, and embeddings were generated using the SimSiam network trained on simulated subtomograms from CryoETSim. The resulting vectors were projected into a 2D UMAP space for visualization.

Mapping the size of each instance, estimated from its voxel count (volume), revealed a clear gradient in the UMAP space, indicating that the learned embeddings encode geometric variation directly from experimental data (Extended Data Fig. 4a). To further test whether the embedding space captures structural similarity, we embedded simulated subtomograms of the 70S ribosome, its 50S and 30S subunits, and RNA polymerase from CryoETSim and mapped them into the experimental space (Fig. 6a). Visualization of the individual instance masks confirmed that these clusters correspond to meaningful macromolecular structures. For *C. reinhardtii*, the simulated 70S ribosomes overlapped with particles resembling eukaryotic 80S ribosomes, consistent with their shared morphology at this resolution. Together, these results show that CryoSiam captures both size- and shape-dependent features of macromolecular complexes, providing a unified embedding space for exploring tomographic information across simulated and experimental domains.

**Figure 6:**
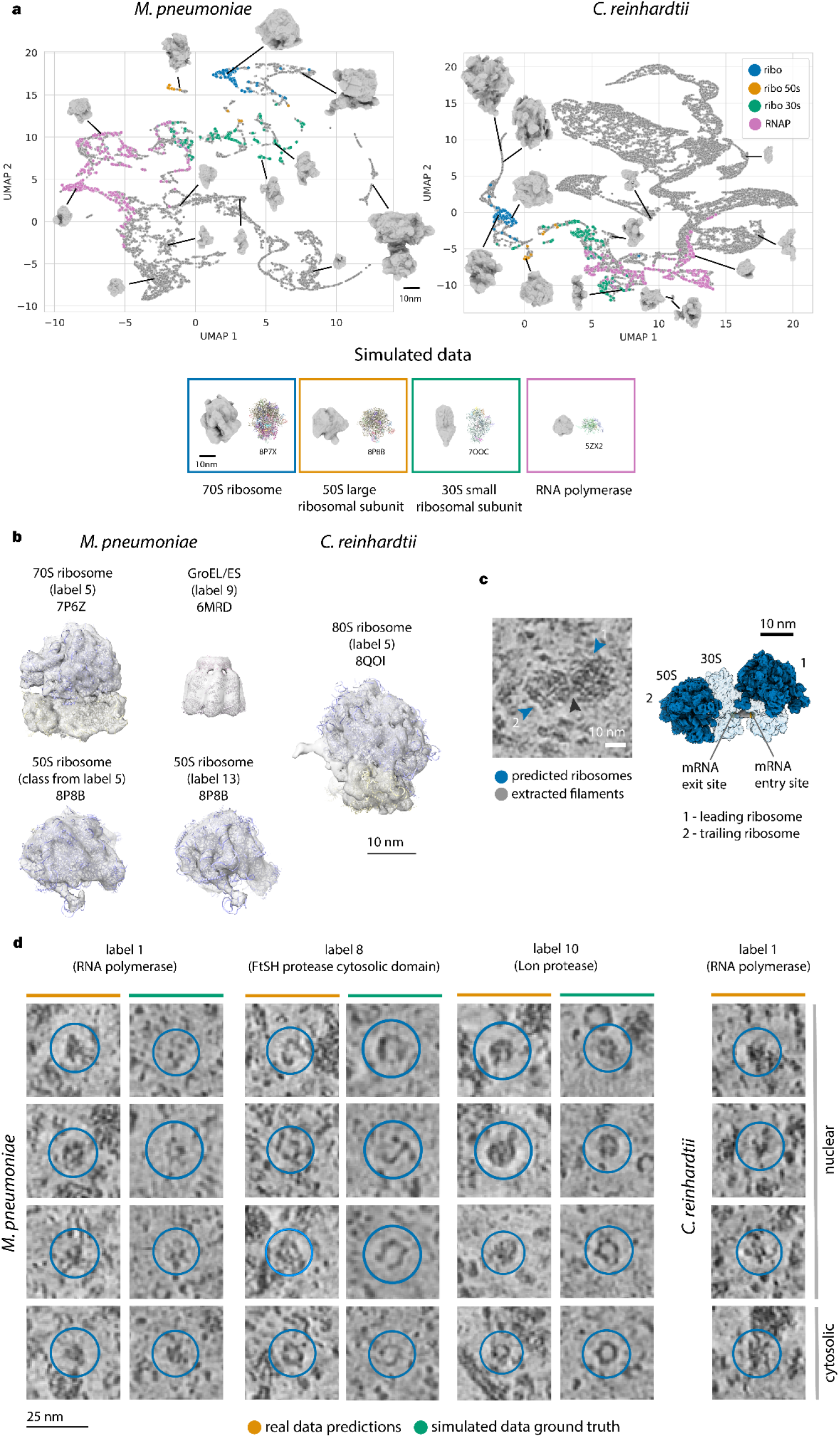
Molecular identification and structural interpretation of CryoSiam predictions in experimental tomograms. **a**. UMAP visualization of subtomogram embeddings from CryoSiam instance predictions in *M. pneumoniae* and *C. reinhardtii*. Each point represents an embedded particle instance, extracted from the denoised tomograms using the corresponding instance masks. The displayed subtomogram volumes from these denoised regions illustrate what individual points in the embedding space correspond to. Reference embeddings from simulated CryoETSim minimal bacterium tomograms (70S ribosome, 50S and 30S ribosomal subunits, and RNA polymerase) are projected into the same space, showing clear separation between molecular classes and similarity in shape and size with the particle instances from the real data. **b**. Subtomogram averaging of CryoSiam-predicted particles from *M. pneumoniae* and *C. reinhardtii*. In *M. pneumoniae*, one of the 50S averages was obtained as a subclass derived from the 70S ribosome picks. Atomic models are fitted (PDB entry indicated) demonstrating consistency with published structures. **c**. Example of a CryoSiam-predicted mRNA filament (grey) bridging two ribosomes (blue), extending from the mRNA exit site of the trailing ribosome (2) to the mRNA entry site of the leading ribosome (1) in *M. pneumoniae*. Ribosomes are shown as the subtomogram average backprojected into the tomogram. **d**. Representative tomographic slices showing CryoSiam predictions on real tomograms (orange) compared with their simulated counterparts from CryoETSim (green). In *C. reinhardtii*, RNA polymerase (label 1) was detected in both nuclear and cytosolic regions, with examples shown for each, where the cytosolic pick is a clear false positive.

To evaluate CryoSiam’s capability for particle identification, we adapted its semantic segmentation task to predict molecular identities as categorical voxel-level classes, focusing on the *M. pneumoniae* tomograms. Because fully unsupervised clustering would complicate evaluation and increase false positives, we employed a semi-supervised strategy using simulated reference data. Fourteen macromolecular complexes were selected for simulation (Extended Data Fig. 4b), including one (label 3) absent in *M. pneumoniae* as a negative control. A new dataset, minimal bacterium v2, was generated by enriching these complexes with structures from the general CryoETSim sample (Supplementary Table 1). During training, both the encoder and decoder were unfrozen and optimized jointly, enabling full network adaptation to this more complex identification task. The resulting predictions were evaluated by subtomogram averaging to assess structural consistency and correspondence with known molecular species.

Subtomogram averaging (STA) was performed using the predicted coordinates (Methods, Fig. 6b). Because only a small number of predictions were obtained for several classes across the 65 tomograms, we performed STA only for labels with more than 100 predicted picks in *M. pneumoniae*, which excluded labels 2, 3, 6, and 12. We carried out two consecutive STA rounds: the first produced averages whose size and morphology aligned with the expected macromolecular targets (Extended Data Fig. 4c,d). In the second STA round, we proceeded with the 70S ribosome (label 5), 50S ribosomal subunit (label 13), and the GroEL ring (label 9) in *M. pneumoniae*, as their initial maps displayed defined features consistent with previously reported structures. Notably, although label 9 was intended to target the GroEL complex, the resulting average corresponded instead to GroES-bound GroEL, likely reflecting the predominant species present *in situ*. These maps were further refined (Methods, Supplementary table 13), resulting in high quality subtomogram averages (Supplementary Fig. 4) that aligned well to atomic models (Fig. 6b). While previous studies have used the *M. pneumoniae* (EMPIAR-10499) dataset focused primarily on the 70S ribosome for particle localization and subtomogram averaging^16–18,42,43^, CryoSiam additionally recovered high-quality averages of the 50S ribosomal subunit and GroEL/ES complex, extending structural characterization beyond what has been previously demonstrated.

Although the model was not trained on eukaryotic ribosomes, predictions in *C. reinhardtii* nonetheless revealed densities that appeared to correspond to 80S ribosomes when using the label associated with the 70S ribosome (label 5; Extended Data Fig. 4c,d and Fig. 6b). Applying the same STA pipeline yielded a high-quality subtomogram average (Supplementary Table 13) that is consistent with the atomic model of the 80S ribosome (Fig. 6b).

Mapping the *M. pneumoniae* ribosome average back into the tomogram revealed regions of active translation, where multiple ribosomes arranged along a continuous mRNA. The mRNA was segmented as a separate instance particle (Fig. 6c), enabling direct visualization of its path between the leading and trailing ribosome.

Few additional complexes were detected in the publicly-available *M. pneumoniae* dataset, but not in sufficient numbers for STA. Therefore, we assessed CryoSiam’s predictions qualitatively (Fig. 6d). Representative examples show a strong visual correspondence between experimental predictions and simulated densities, consistent in size and morphology. These observations support the structural plausibility of the predictions, although a larger dataset will be required to confirm full molecular identities with STA. Notably, for *C. reinhardtii* CryoSiam also detected densities resembling RNA polymerase in cytoplasmic regions, underscoring the need for careful interpretation when extending predictions beyond validated targets.

## Discussion

Automated computational approaches are essential for extracting meaningful information from the rapidly growing volume of cryo-ET data^9,11^. However, the lack of voxel-level ground truth and the strong variability in noise, contrast, and structures across datasets have long limited the use of deep learning in cryo-ET^21,44^. While simulated data offer a practical alternative for training and evaluation^3,13,23^, differences from experimental data in terms of realistic representation of modulations arising from electron imaging and complexity of biological systems have hindered direct transfer of trained models^7,45,46^.

CryoSiam bridges this gap by combining a physically realistic simulated dataset, CryoETSim, with a self-supervised learning framework tailored to the statistical and physical properties of cryo-ET. Through carefully designed transformations that emulate defocus variation, frequency modulation, and noise, CryoSiam learns representations with self-supervised learning that generalize across simulated and experimental tomograms. This approach extends the recent progress of self-supervised frameworks such as TomoTwin^16^, CryoSAM^18^, and MiLoPYP^17^, demonstrating that high-quality representations learned from simulated data can be directly transferred to real data.

At the voxel level, CryoSiam performs denoising, semantic segmentation, and instance segmentation directly on experimental data without retraining. While existing denoising approaches such as Cryo-CARE^40^, IsoNet^47^, Topaz-Denoise^48^, rely on dataset-specific training, CryoSiam achieves effective denoising on real data to suppress noise and restore contrast by directly applying models that have been trained through supervised learning on clean simulated tomograms. The semantic segmentation task further distinguishes membranes from surrounding particles, enabling rapid identification of membrane-associated complexes without additional post-processing or supervised models. Complementing this, the instance segmentation task isolates individual macromolecular complexes by masking surrounding noise and separating overlapping densities. This capability is particularly valuable for downstream structural analysis, as it provides accurate particle-level masks that can be directly used for subtomogram extraction and embedding generation, even when molecular identity is unknown. In practice, accurate particle masks were the key success factor for subsequent particle identification, highlighting the critical role of instance segmentation within the CryoSiam workflow. Trained entirely on simulated data, all of the three tasks transfer robustly to experimental tomograms, providing a practical framework for exploring cellular architecture and accelerating structural discovery within complex biological environments.

At the subtomogram level, CryoSiam extends standard self-supervised embeddings by linking them to instance segmentation masks and denoised tomograms, allowing each embedded point to be traced back to its corresponding particle in the experimental data and directly visualized. In simulated data, where particle masks are accurately defined, CryoSiam reliably distinguished macromolecular complexes across a wide molecular-weight range, demonstrating its capacity to capture fine structural differences even among smaller particles. When applied to experimental data from *M. pneumoniae*^*37*,38^ and *C. reinhardtii*^*39*^, embeddings derived from simulated training generalized effectively, grouping structurally related complexes and linking 70S and 80S ribosomes across species.

Using a semi-supervised particle identification strategy with simulated references, CryoSiam further enabled automated detection of molecular complexes, with subtomogram averaging validating the identities of fully assembled eukaryotic and prokaryotic ribosomes, 50S large subunit, and GroEL/ES directly from experimental predictions. However, verifying particle identification still relies on subtomogram averaging, which remains sensitive to particle number, structural and conformational heterogeneity, and signal-to-noise ratio. When predicted instances are few or the macromolecular complex is small, averaging often fails to converge to meaningful structures, limiting systematic validation of automated detections. Thus, while CryoSiam advances the field toward data-driven particle identification, the challenge of reliable confirmation at low counts remains an open problem for future methodological development. Future work will integrate validated experimental picks to retrain CryoSiam on combined simulated and real tomograms, advancing toward a supervised model that better captures structural and conformational diversity.

Together, these results show that CryoSiam bridges the long-standing gap between simulated and experimental cryo-ET data, capturing biologically meaningful representations from local voxel features to whole macromolecular complexes. By providing pretrained models and a large-scale simulated dataset, CryoSiam establishes a foundation for transferable and automated analysis of tomograms, paving the way for systematic, in situ mapping of macromolecular organization.

## Methods

### Simulated data generation

Tomographic datasets were simulated using CTS^21^, generating a total of 400 tomograms across three biologically distinct sample types. The general sample (300 tomograms) contained 141 PDB-derived structures together with membranous vesicles, membrane-embedded complexes, smaller cytosolic complexes (distractors), actin filaments, and microtubules. The minimal bacterium sample (50 tomograms) included complete 70S ribosomes, ribosomal subunits, bacterial RNA polymerase, membranous vesicles, membrane-embedded complexes, distractors, and simulated DNA filaments. The chromatin sample (50 tomograms) consisted of various nucleosome structures and DNA filaments. Molecular weights in the general sample ranged from 10 kDa to >4000 kDa (Supplementary Table 2). To model crowded cellular environments, particles were sequentially placed from largest to smallest with the brute force model filling^21^, achieving crowding densities of up to 80%. Sample thicknesses ranged from 60-100 nm for the general sample and 35-100 nm for the minimal bacterium and chromatin samples.

DNA filaments in the chromatin and minimal bacterium samples were simulated using the PolymerCpp software^49^, which implements the wormlike-chain (WLC) model^32^. Filaments were generated with path lengths corresponding to short DNA segments of 50, 100, 150, 200, 250, 300, 350, 400, 450, 500, and 1000 base pairs. The persistence length, defining the stiffness of the polymer chain, was sampled between 5 and 20 base pairs (5, 6, 8, 10, 20). These parameters were selected to capture variability in DNA folding and compaction in the simulations. The resulting 3D conformations were converted into PDB files and incorporated into the CTS simulations alongside other macromolecular components (Supplementary Fig. 1). The RNA filaments were separated from the existing PDB entries (transcription/translation complexes) that contained the macromolecules interacting with RNA.

Each tomogram was generated by projecting a 3D structural model into a synthetic tilt series from −60° to +60° in 2° increments. Imaging parameters included a defocus range of 2-5 µm, pixel size of 6.8 Å/pixel, acceleration voltage of 300 kV, amplitude contrast of 2.7, Gaussian noise (σ = 0.2), and a total electron dose of 120 e−/Å−^2^. Each projection was corrupted with additive noise to generate the realistic tilt series and after that modulated by a contrast transfer function (CTF) corresponding to the selected defocus. Both the clean and CTF-modulated/noisy tilt series were then independently reconstructed into 3D tomograms using weighted back-projection (WBP) implemented in IMOD^50^.

To support downstream deep learning applications, the original CTS code was modified to additionally export instance-level annotations, including per-particle instance masks, spatial coordinates, and volume measurements. The final CryoETSim dataset, therefore, contains clean and noisy tilt series, CTF-modulated tomograms, voxel-wise semantic masks for macromolecular identities (background, membranes, microtubules, actin filaments, DNA/RNA filaments, and particle classes), and instance masks with corresponding coordinate and volume metadata. Together, these outputs provide a comprehensive and realistic resource for training self-supervised models and benchmarking segmentation or detection methods in cryo-electron tomography.

### SimSiam self-supervised network architecture

SimSiam takes two views, *x*_1_ and *x*_2_, of the subtomogram *x* (Fig. 1c). Different random augmentations are applied independently as transformations *T*_1_ and *T*_2_ to generate these views. The augmented views, *x*_1_ and *x*_2_, are then passed through an encoder, based on a ResNet architecture^30^, to extract meaningful features. A multilayer perceptron (MLP) head projects the encoded features into a representation space, producing global embeddings 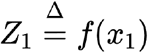 and 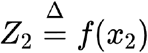, where the encoder parameters are shared between both views.

A predictor *h* is then used to generate modified representations 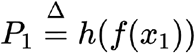 and 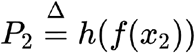. The optimization objective minimizes the negative cosine similarity between the two representations, defined as:

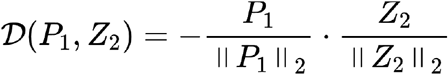

To prevent representation collapse^27^, the predictor *h* and a stop-gradient operation are employed while minimizing the negative cosine similarity:

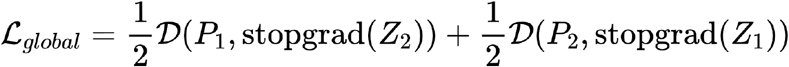

### DenseSimSiam self-supervised network architecture

DenseSimSiam extends SimSiam by incorporating voxel-wise dense representations. The first part of the architecture follows the SimSiam structure, while the second part introduces a decoder, based on a Feature Pyramid Network (FPN)^31^, with a 1 × 1 3-layer CNN projector at its final layer (Fig. 1b). The decoder generates dense voxel-level embedding representations 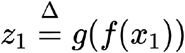 and 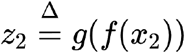, where the weights are shared across views. A 1 × 1 2-layer CNN predictor *h*^′^ then produces modified dense representations 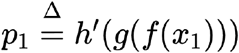 and 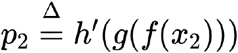. The cosine similarity loss for the dense embeddings is defined as:

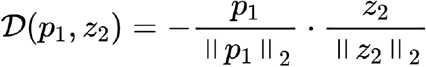

To prevent collapsing while learning dense embeddings, the predictor *h*^′^ and the stop-gradient operation are used. The loss function is computed over voxels in the overlapping regions of *x*_1_ and *x*_2_:

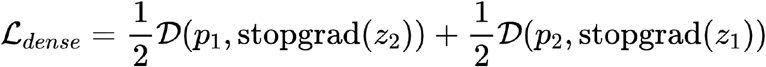

To capture multi-scale representations, a 1 × 1 3-layer CNN projector is applied to different levels of the decoder, producing embeddings at multiple resolutions:

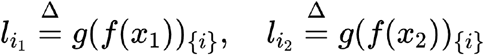

where *i* denotes the -th level of the decoder. A 1 × 1 2-layer CNN predictor *h*^′^ generates the corresponding modified representations:

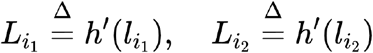

The cosine similarity loss is then computed for each level *i*:

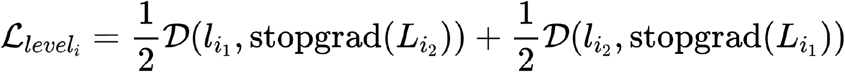

The total level-wise loss is obtained by summing the individual losses from all *n* levels:

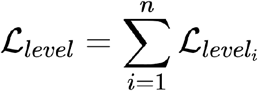

The final training loss combines the global, dense, and multi-scale losses:

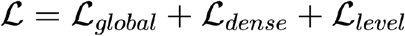

### Image Transformations

#### Random Gaussian Noise

To enhance robustness against noise, random Gaussian noise is introduced by adding Gaussian-distributed values (sigma: 0.1-0.5) to voxel intensities. This transformation simulates the high-noise levels in low-dose cryo-ET data, ensuring that embeddings focus on structural features rather than noise artifacts.

#### Random Gaussian Low-Pass Filtering

To account for variations in resolution, low-pass filtering attenuates high-frequency details while preserving coarse structural features. A Gaussian filter with a randomly selected sigma (0.5-2) smooths voxel intensities, reducing sensitivity to resolution differences and ensuring consistency across datasets. This transformation approximates the effects of imaging at varying defocus levels, where higher defocus corresponds to stronger low-frequency emphasis in the recorded signal.

#### Random Gaussian High-Pass Filtering

High-pass filtering enhances fine structural details by attenuating low-frequency background components while preserving high-frequency information. The transformation is implemented by subtracting a more strongly smoothed version of the tomogram (σ = 1-1.5) from a lightly smoothed version (σ = 0.1-0.5), which effectively acts as a high-pass filter. This transformation approximates the contrast conditions observed at lower defocus values, where higher-frequency information is better preserved in the recorded images.

#### Masking-Out Voxels Transformation

To encourage context-aware feature extraction, masking-out is applied, where 10-50% of voxels are randomly removed per view. The model learns to infer missing voxel information by leveraging structural context, enforcing consistency between masked and unmasked regions across different views.

#### Subtomogram Affine and Cropping Transformations

Affine transformations introduce spatial variability through random rotations (0-360°) and translations (up to 10 voxels) in any direction, simulating orientation and positional changes. Additionally, random cropping removes up to 20% of the subtomogram, enhancing model robustness by ensuring invariance to positional and orientation-based differences.

### Contrastive learning

Pairs of subtomograms are passed through the shared SimSiam-pretrained backbone to generate normalized embeddings *e*_1_ and *e*_2_. A binary label is assigned, where *y* = 1 for positive pairs (same class) and *y* = 0 for negative pairs (different classes). The contrastive loss is defined as:

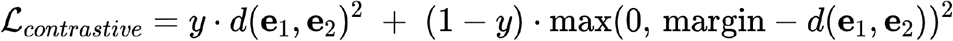

where the distance between embeddings is computed as:

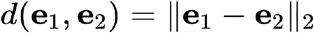

Positive pairs are pulled closer together by minimizing *d*(*e*_1_, *e*_1_) while negative pairs are pushed apart by enforcing a minimal separation margin = 1. This contrastive refinement enhances the discriminative power of the learned embeddings, improving class separability with minimal labeled data.

### Downstream tasks

#### Denoising

Denoising reduces the high noise levels inherent in cryo-ET while preserving structural features. The Mean Squared Error (MSE) loss is used to optimize performance:

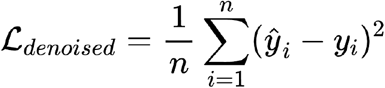

where *ŷ*_*i*_ is the predicted voxel intensity, *y*_*i*_ is the ground truth, and is the number of voxels.

#### Semantic segmentation

Semantic segmentation classifies each voxel into predefined structural classes and predicts distances from boundaries. The loss function for this task consists of three components. The first is the Cross-Entropy Loss, ℒ_*sem*1_, which ensures accurate voxel-wise classification:

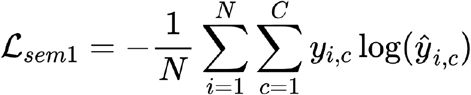

where *N* is the number of voxels, is the number of classes, *y*_*i,c*_ is the ground truth probability for class *c* at voxel *i*, and *ŷ*_*i,c*_ is the predicted probability.

To address class imbalance, the Generalized Dice Loss, ℒ_*sem*2_, is incorporated:

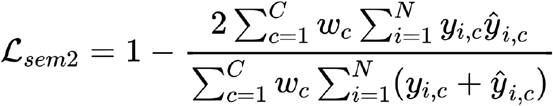

where 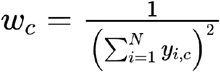 ensures proper weighting of each class.

In addition, the Mean Squared Error (MSE) loss, ℒ_*distance*_, is applied to refine the predicted distances from the background:

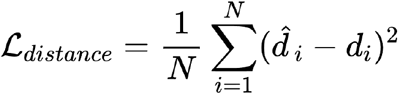

where 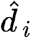 and *d*_*i*_ are the predicted and ground truth distances.

The final loss function for semantic segmentation is defined as:

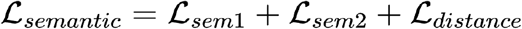

#### Instance segmentation

Instance segmentation predicts particle boundaries, foreground regions, and distances from the background using a watershed segmentation algorithm^33^, refined with a multi-cut algorithm^35^. The loss function for this task consists of three terms. The first, ℒ_*distance*_, is an MSE loss that optimizes the predicted distances of voxels from the background:

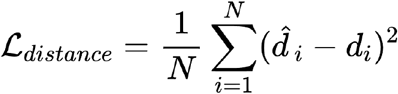

The second term, ℒ_*boundary*_, is a Binary Cross-Entropy (BCE) loss applied to the predicted boundaries:

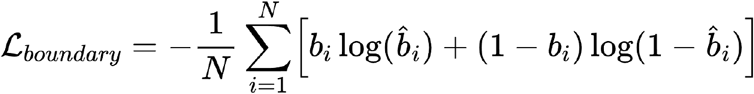

where *b*_*i*_ is the ground truth boundary label.

The third term, ℒ_*foreground*_, is another BCE loss used to classify foreground voxels:

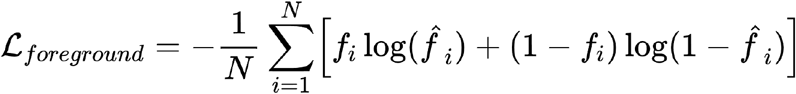

where *f*_*i*_ is the ground truth foreground label.

The final instance segmentation loss function is defined as:

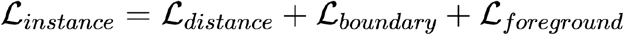

#### Particle subtomogram classification

Particle classification was performed using k-nearest neighbors (k-NN) and a multilayer perceptron (MLP). For k-NN, the cosine distance was applied when evaluating embeddings from SimSiam without contrastive learning, while the Euclidean distance was applied to contrastive learning-adapted embeddings. The classification results were reported with *k* = 1.

The MLP classifier was trained using the categorical cross-entropy loss:

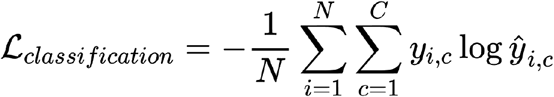

where *y*_*i,c*_ represents the one-hot encoded ground truth class label for the subtomogram *ŷ*_*i,c*_, and is the predicted probability for class.

### Training strategy

The training protocol consisted of self-supervised pretraining followed by task-specific training for semantic segmentation, instance segmentation, particle identification, and contrastive learning.

DenseSimSiam was trained using cosine similarity as the loss function for 600 epochs with a cosine annealing scheduler^51^, an initial warmup of 5 epochs, and a maximum learning rate of 0.5. Optimization was performed using Stochastic Gradient Descent (SGD) with a momentum of 0.9, a weight decay of 0.00001^52^, and a batch size of 10.

SimSiam followed a similar strategy, trained for 200 epochs with a batch size of 10, using a cosine annealing scheduler with a 10 epoch warmup and a maximum learning rate of 0.05. Optimization was performed with SGD, using a momentum of 0.9 and a weight decay of 0.00001.

For downstream tasks and contrastive learning, a one-cycle learning rate scheduler^53^ was used, with a fixed learning rate of 0.001 and a training duration of 200 epochs. Batch sizes varied per task. The AdamW optimizer^54^ was used for stability and regularization.

### Evaluation metrics

The performance of the CryoSiam framework was assessed using specialized metrics for semantic segmentation, instance segmentation, and particle identification.

For semantic segmentation, the Sørensen-Dice coefficient (Dice) measured the overlap between predicted and ground truth masks:

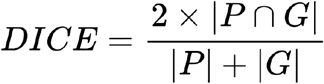

where *P* and *G* represent the predicted and ground truth sets, respectively. A Dice score of 1 indicates perfect overlap, while 0 signifies no overlap.

For the instance segmentation, the Average Precision (AP) metric evaluated the model’s ability to localize and separate particle instances. The Intersection over Union (IoU) was calculated as:

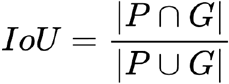

where *P* and *G* are the predicted and ground truth masks. A prediction was considered correct if the IoU exceeded a predefined threshold, such as 0.5 (AP-50). Precision was computed as:

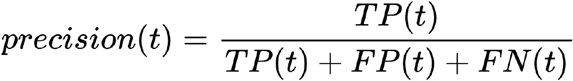

where *TP, FP*, and *FN* represent true positives, false positives, and false negatives at threshold. The overall AP was defined as:

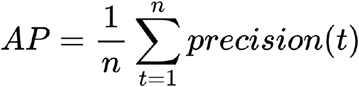

where *n* represents the total number of thresholds considered.

For particle identification, the F1 score evaluated classification and localization accuracy:

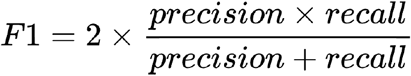

where precision and recall were given by:

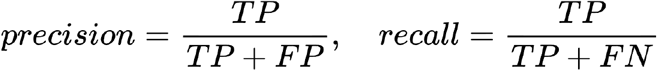

A high F1 score indicates a balance between identifying relevant particles (recall) and minimizing false detections (precision).

### Ablation studies

Ablation studies were conducted using simulated tomograms from the CryoETSim dataset to evaluate the impact of core architectural and training choices within the CryoSiam framework (Supplementary Tables 6-11). By independently varying data augmentations, embedding dimensions, and loss components, we quantified how each factor influenced the quality of voxel-level embeddings and their ability to directly support semantic classification. The findings informed the parameter settings used in the final trained model.

To assess the quality of the learned embeddings directly, a simple three-layer CNN head with 1×1 convolutions was trained on top of a frozen encoder to classify voxels into six semantic categories: background, membrane, particles, microtubules, actin, and DNA filaments. The minimal design of this head was chosen to avoid introducing additional spatial context or complex learned functions, ensuring that performance reflected only the representational quality of the embeddings. The model was trained on a reduced dataset comprising 40 general, 10 minimal bacterium, and 10 chromatin tomograms, with 80 percent used for training and 20 percent for testing (Supplementary Table 6). Performance was evaluated using the Dice score. The analysis assessed the effects of different augmentations, including Gaussian noise, low-pass and high-pass filtering (Supplementary Table 7), voxel masking (Supplementary Table 8), and embedding dimensionality for voxel-level (Supplementary Table 9), global level (Supplementary Table 10), and the incorporation of multiple levels of embeddings (Supplementary Table 11).

### Experimental cryo-ET data

All experimental tomograms used in this study were obtained from publicly available datasets deposited in the Electron Microscopy Public Image Archive (EMPIAR)^55^ and the Chan Zuckerberg Imaging Institute (CZII) CryoET Data Portal^56^. Specifically, cellular cryo-electron tomograms of *M. pneumoniae* (EMPIAR-10499, DS-10003) and *C. reinhardtii* (EMPIAR-11756, DS-10301) were used for predictions on experimental data (Supplementary Table 12).

For *M. pneumoniae*, predictions were performed directly on the tomograms available through the CryoET Data Portal, which had been preprocessed using Warp and refined in M as described in the original publication^37^. For *C. reinhardtii*, the motion-corrected tilts and tilt alignments were downloaded from EMPIAR and imported into Warp-1.0.9^57^ for 2D and 3D CTF-estimation, and reconstruction of tomograms and subtomograms. To minimize differences in preprocessing that might influence model performance, we performed template matching of ribosomes using PyTOM^58^, followed by removal of false positives using RELION 3D classification^59^, and finally image-warp, stage-angle and defocus correction in M-1.0.9^37^. The refined tomograms were then reconstructed in Warp and used for CryoSiam predictions.

For both datasets, the corresponding membrane segmentation predictions and the *C. reinhardtii* Cryo-CARE denoised tomograms were retrieved from the CryoET Data Portal to facilitate direct comparison with CryoSiam predictions. All experimental tomograms were used without additional normalization or filtering beyond the procedures described above.

### CryoSiam predictions on experimental cryo-ET data

CryoSiam voxel-level downstream tasks, tomogram denoising, lamella mask prediction, semantic segmentation, and instance segmentation, were applied directly to experimental tomograms of *M. pneumoniae* (EMPIAR-10499) and *C. reinhardtii* (EMPIAR-11756), reconstructed at voxel level of 6.8 Å/pixel and 7.84 Å/pixel respectively. All models were trained on simulated tomograms from the CryoETSim dataset and used without additional fine-tuning.

For denoising, experimental tomograms were scaled (clipping values of 0.01 and 99.9 percentiles from the intensity), inverted in contrast and the denoised tomogram was predicted with the CryoSiam denoising network. For qualitative comparison, Gaussian-filtered tomograms were generated using a standard 3D Gaussian kernel, and corresponding Cryo-CARE denoised tomograms were obtained from the CryoET Data Portal^56^ for the *C. reinhardtii* dataset (DS-10301).

A lamella mask prediction network was trained on denoised simulated tomograms with binary labels distinguishing cellular/lamella regions from surrounding regions. The resulting masks were applied to experimental tomograms to isolate the physical sample volume and exclude background during subsequent analyses. Semantic and instance segmentation networks were then applied to the denoised and masked tomograms. For integrated visualization, semantic membrane masks and instance predictions were combined to identify macromolecular complexes located within two voxels of membrane surfaces. All results were visualized using ChimeraX^60^ and napari^61^, with each structural component rendered in a distinct color. Voxel-level predictions were performed on overlapping 128^3^-voxel patches, which were subsequently merged to obtain the full tomogram output.

Predicted instances were cropped using convex-hull masks, zero-padded to 64^3^ voxels, and embedded with the SimSiam network trained on simulated subtomograms to obtain representation vectors projected into 2D UMAP space. For particle identification, a semi-supervised strategy was adopted: the network was trained on a new minimal bacterium v2 dataset, containing 80 tomograms with 14 molecular classes as target for the particle identification, generated in the same strategy as the general CryoETSim tomograms. The encoder, decoder, and segmentation heads were jointly optimized from self-supervised pretrained weights using the semantic segmentation training strategy. Predicted particle centroids were exported in a STAR format for subtomogram averaging.

### Subtomogram averaging

To evaluate the quality of CryoSiam particle identification, we performed subtomogram averaging (STA) using the predicted particle coordinates as input. Initial reference maps were generated at 40 Å resolution from the corresponding PDB structures in ChimeraX^60^, resampled to match the pixel size of each tomogram in bin 4 (6.8 Å/pixel for EMPIAR-10499 and 7.8 Å/pixel for EMPIAR-11756). Subtomograms were aligned to these references using SUSAN, a cross-correlation-based STA framework^62,63^. Alignment was carried out through five iterations covering the full angular search space, followed by five additional iterations restricted to a smaller angular range.

Once the particles were centered with the alignment, the resulting coordinates and orientations were exported as STAR file format compatible with Warp-1.0.9^57^ for subtomogram extraction. RELION-4.0.1^59^ was then used to perform multi-class ab initio model generation.

For the first round of STA, only particles contributing to well-resolved ab initio classes were retained for further refinement. The selected particles and resulting ab initio model were used to initialize a second SUSAN project, employing a box size of 96 pixels, a fixed bandpass of 30 Å, and data limited to the central tilt range (±30°, not adjusted for lamella pretilt). Five iterations of global alignment were followed by five iterations of local refinement, after which a per-tilt 2D shift refinement was performed for 20 iterations. Final maps were reconstructed after discarding outlier particles based on their cross-correlation (CC) score. In cases where the CC distribution was multimodal, a multi-reference alignment was applied instead. This STA pipeline was applied to all analyzed target proteins. For label 9 and label 14, C7 symmetry was imposed during the second SUSAN refinement to compensate for the limited number of particles.

For the species giving rise to maps with higher resolution features, we performed a second STA round in RELION-4.0.1^59^. The ab initio models previously generated were used to run 3D classification using default parameters with four classes, and particles resembling the molecule of interest were selected for a final round of refinement. For label 5 (ribosome), a 50S ribosomal subunit class was classified out and was also refined separately. For the GroEL/ES complex (label 9), the refinement of the particle symmetry axis was aligned to the Z-axis using relion_align_symmetry using C20 symmetry. A second round of refinement was performed using C7 symmetry, followed by an additional round of 3D classification using local angular searches with symmetry relaxation. The final particles were selected and used for a final 3D refinement. RELION-4.0.1 postprocessing was used to calculate the FSC curves (Supplementary Fig. 4). Final summary of resolutions and particle numbers are summarized in Supplementary Table 13.

## Declarations

### Data availability

The simulated dataset generated in this study, CryoETSim, will be made publicly available upon publication. All accession numbers for PDB models used in the generation of the simulated dataset are listed in Supplementary Table 2. Experimental cryo-ET data used for evaluation are publicly accessible through the Electron Microscopy Public Image Archive (EMPIAR) under accession numbers EMPIAR-100499 (*M. pneumoniae*) and EMPIAR-11756 (*C. reinhardtii*), and through the CryoET Data Portal (https://cryoetdataportal.czscience.com/) under dataset identifiers DS-10003 (*M. pneumoniae*) and DS-10301 (*C. reinhardtii*).

### Code availability

The source code for CryoSiam is available at https://github.com/frosinastojanovska/cryosiam.

A companion visualization tool for inspecting and analyzing CryoSiam predictions is provided at https://github.com/frosinastojanovska/cryosiam_vis.

Comprehensive documentation, including installation instructions and usage examples, is also available. Pretrained CryoSiam models will be made publicly available along with the GitHub repository.

### Contributions

F.S. conceived the project with input from J.B.Z., J.M. and A.K. F.S. developed the CryoSiam framework and CryoETSim dataset, performed all computational analyses, and wrote the initial manuscript with input from J.B.Z., A.K., J.M., and R.K.J. R.M.S. and R.K.J. contributed to the computational structural analysis. J.B.Z., A.K., J.M. supervised the project and acquired funding. All authors contributed to data interpretation and manuscript writing.

## Acknowledgments

Computational analyses and simulations were performed using the High-Performance Computing (HPC) cluster at EMBL and the NHR@TUD HPC facilities. The authors acknowledge the support of these resources, including technical assistance and computational infrastructure, which were essential for this work. Computing time at NHR@TUD was provided by the Federal Ministry of Education and Research (BMBF) and the participating state governments within the framework of the Joint Science Conference (GWK) for national HPC at universities. The authors thank Jurij Pečar for his valuable technical support with HPC resources and Thomas Hoffmann for the technical support with structural biology software on HPC resources. We thank Matt Swulius for providing the PDB files used in developing the CTS simulator. R.K.J. was supported by the Independent Research Fund Denmark (grant number 0164-00010A). R.M.S. was supported by an ARISE fellowship (European Union’s Horizon 2020 Research and Innovation Programme under the Marie Skłodowska-Curie grant agreement number 945405). This work was supported by EMBL core funding to J.B.Z., J.M., A.K., an EMBL Infection Biology Transversal Theme Synergy grant to J.M., and a Chan Zuckerberg Initiative grant for visual proteomics (2021-234620) to A.K. and J.M.

---

**Extended Data Figure 1:**
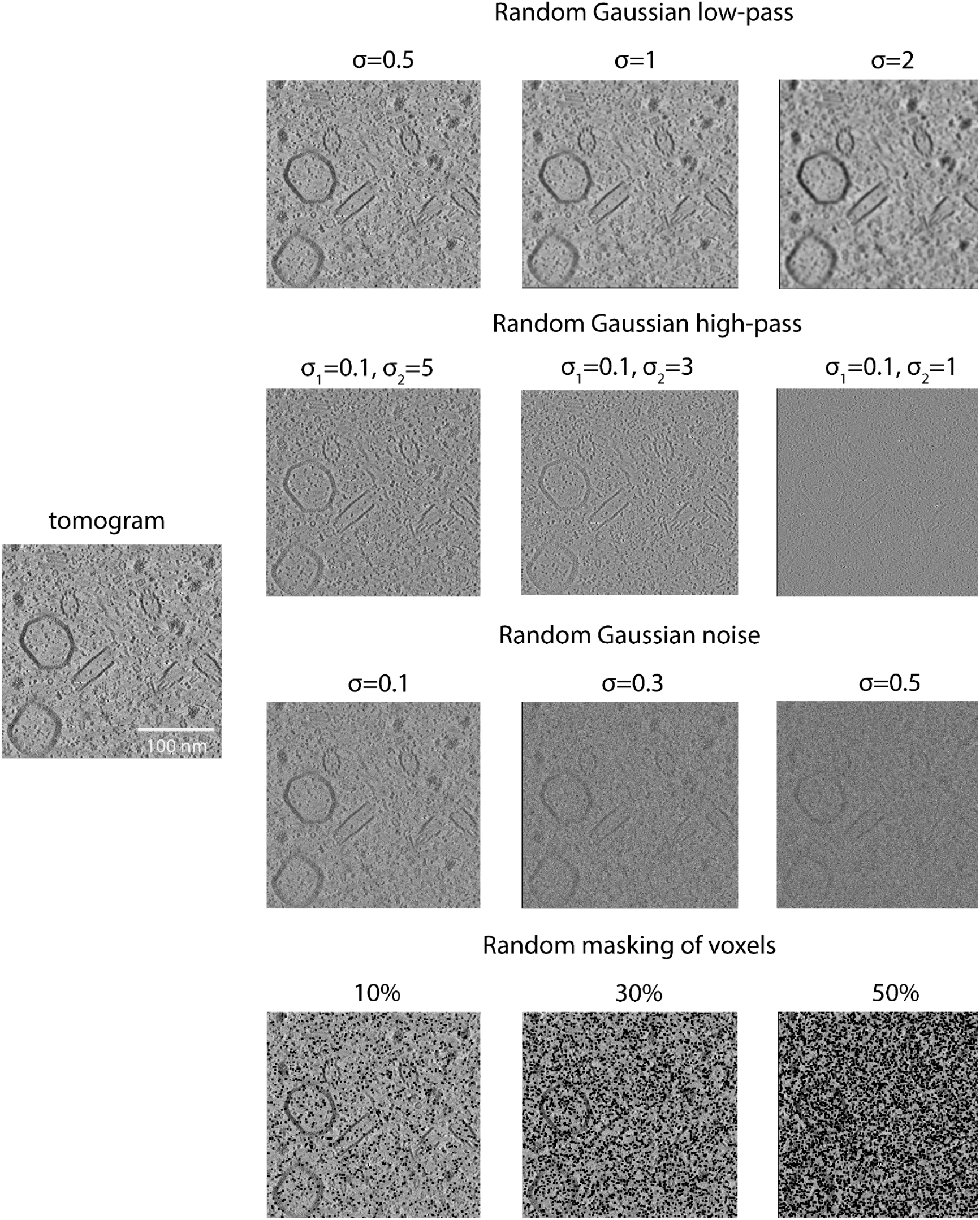
Image transformations used for self-supervised training of the DenseSimSiam module. A representative tomogram (left) and examples of the random image transformations applied during training (right). Transformations include random Gaussian low-pass filtering (σ = 0.5-2), Gaussian high-pass filtering (σ_1_ = 0.1, σ_2_ = 1-5), Gaussian noise addition (σ = 0.1-0.5), and random voxel masking (10-50%). These transformations introduce controlled variation in frequency content, noise level, and missing information to encourage the network to learn invariant and robust voxel-level representations.

**Extended Data Figure 2:**
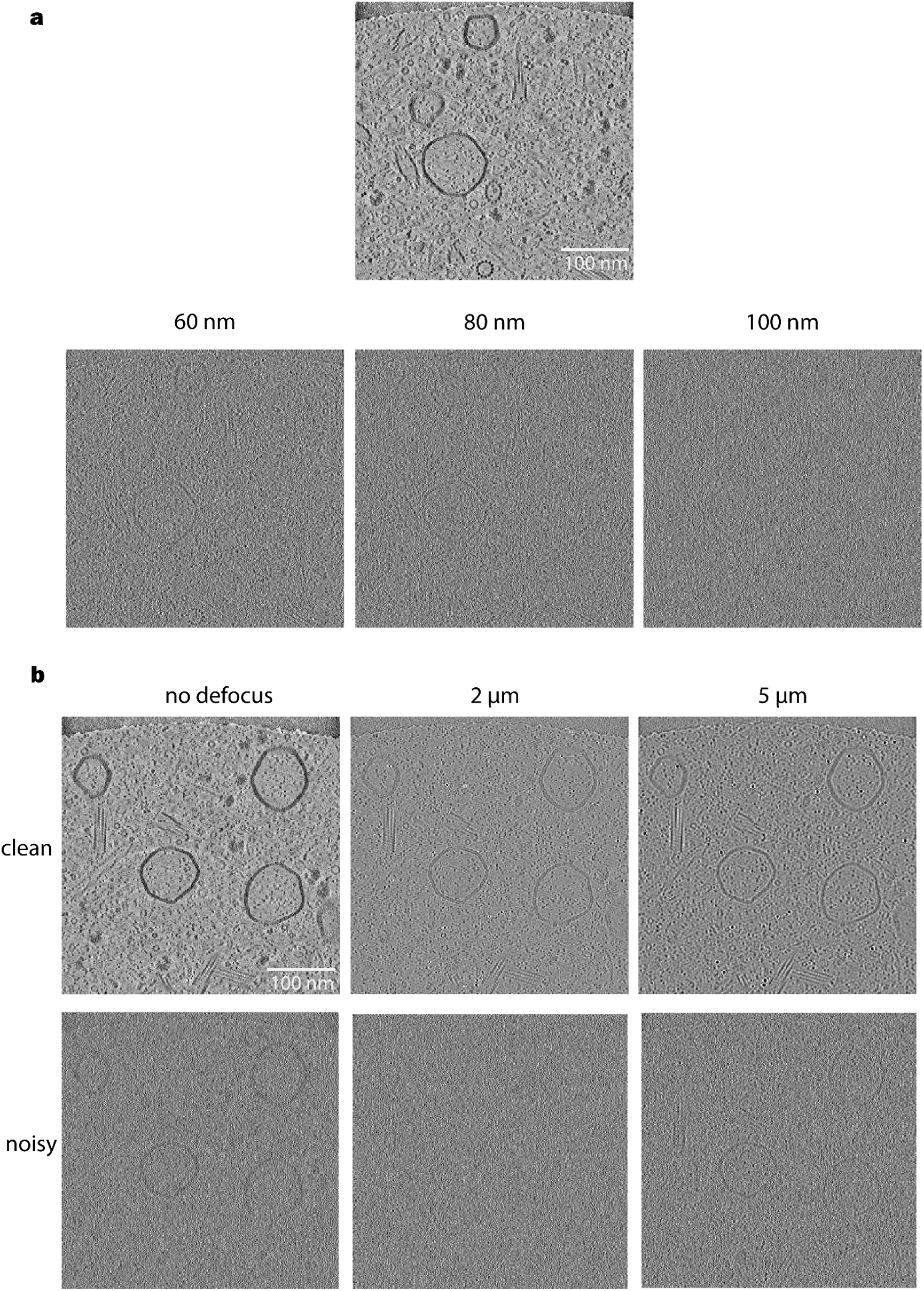
Effects of defocus and sample thickness on contrast and image diversity in simulated tomograms. **a**. Example tomograms simulated at different sample thicknesses (60 nm, 80 nm, and 100 nm). Thinner samples display stronger contrast, while thicker samples exhibit reduced visibility of the information. **b**. Example tomograms simulated with varying defocus values (no defocus, 2 µm, and 5 µm) shown for both clean (top row) and noisy (bottom row) conditions. Increasing defocus enhances low-frequency information and image contrast, whereas lower defocus values preserve high-frequency detail.

**Extended Data Figure 3:**
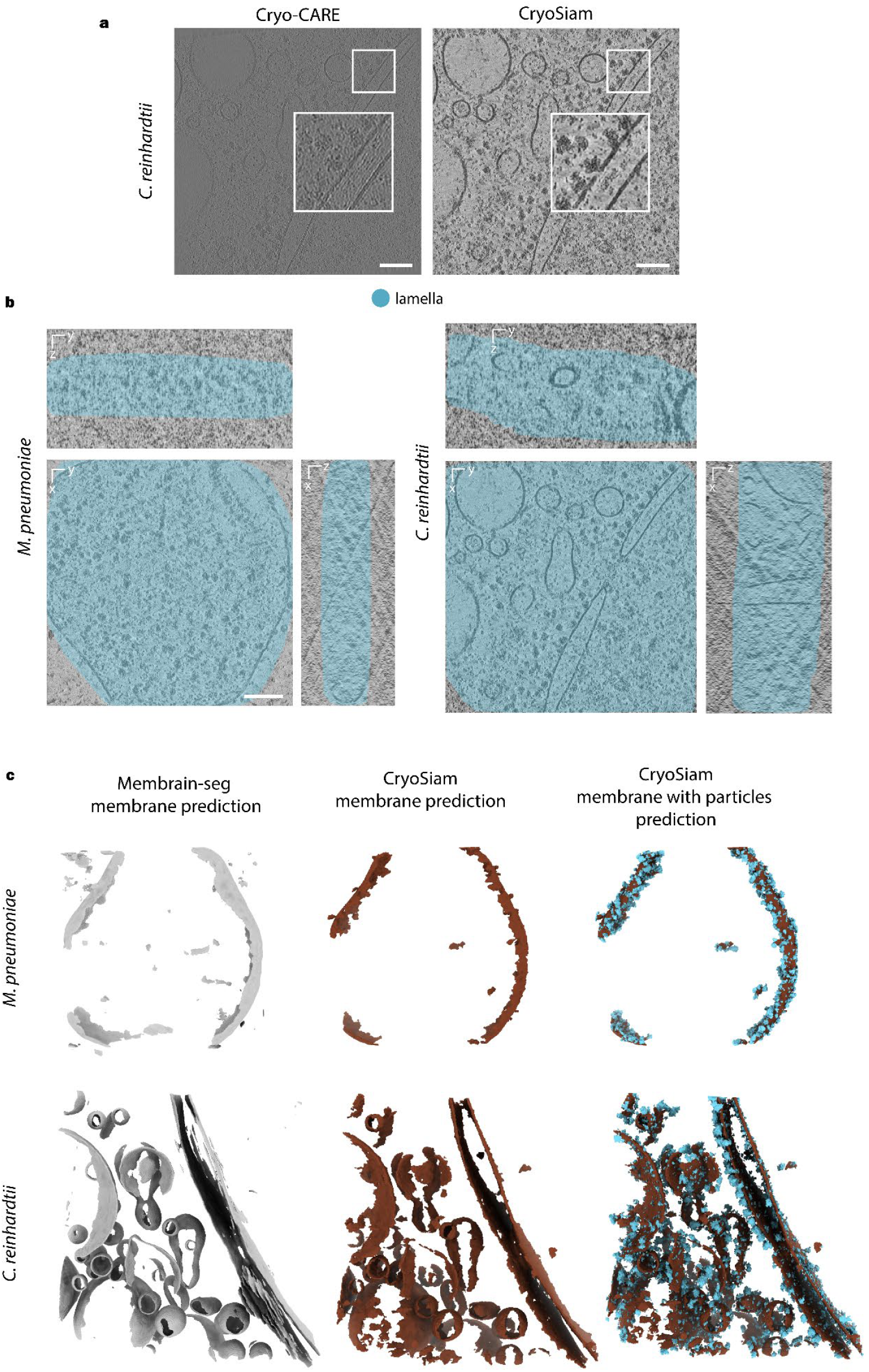
Evaluation of CryoSiam performance on experimental tomograms. **a**. Comparison of denoising methods. CryoSiam denoising compared with Cryo-CARE on a *C. reinhardtii* tomogram. CryoSiam enhances image contrast and recovers fine structural details lost due to defocus modulation. Both tomograms were visualized in napari^61^ without additional intensity scaling. **b**. Cell or Lamella region prediction. A downstream network trained on denoised simulated tomograms predicts the regions containing the sample (cyan) in *M. pneumoniae* (cell) and *C. reinhardtii* (FIB-lamella) data, accurately separating the sample from background noise to reduce false positives in later analyses. **c**. Membrane segmentation comparison. Membrane predictions from the supervised MemBrain-Seg network (left) and CryoSiam (middle) show comparable delineation accuracy. Combined membrane-particle visualizations (right) highlight the added interpretive value of CryoSiam’s multilevel predictions.

**Extended Data Figure 4:**
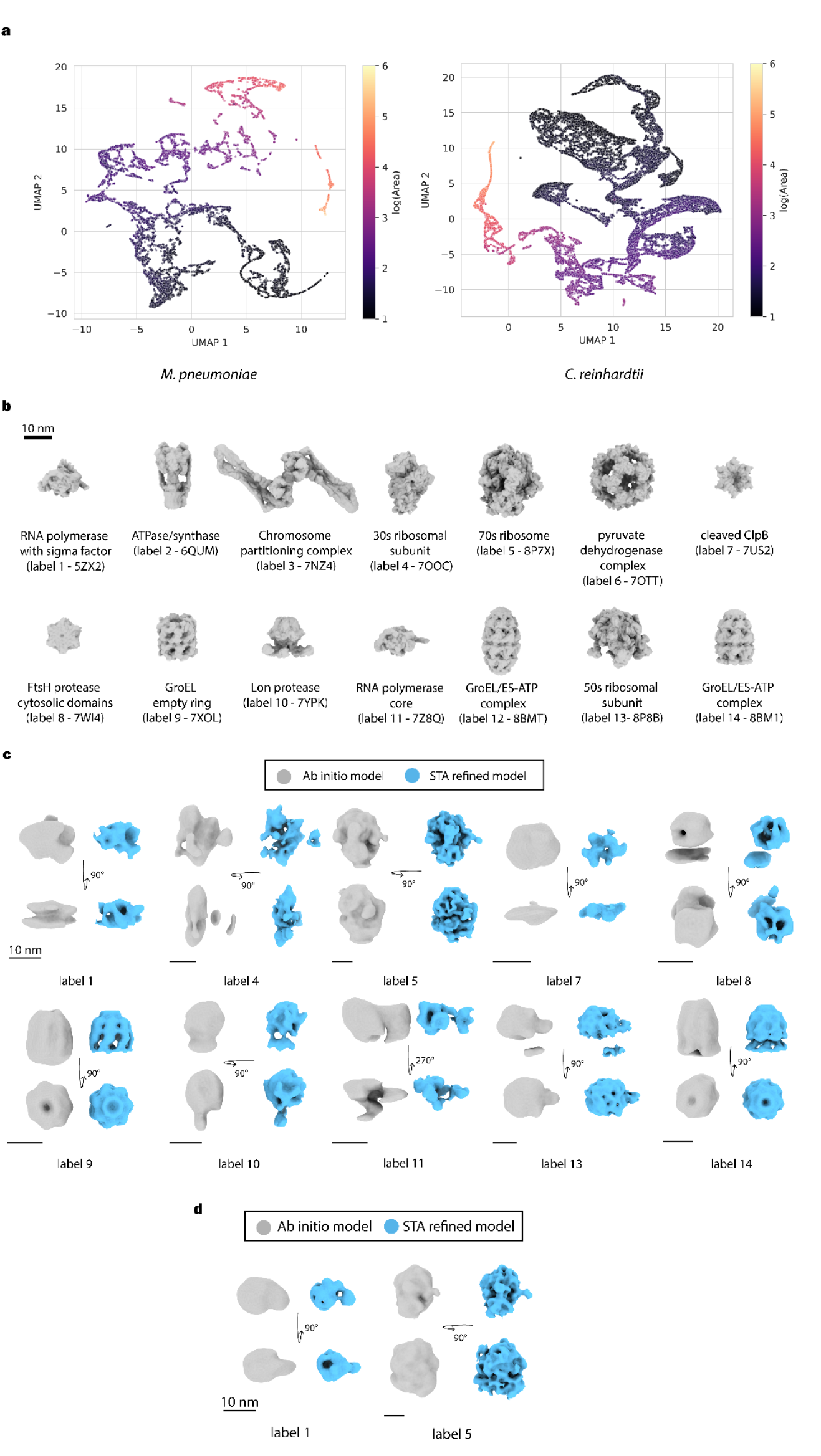
Instance embedding space, simulated training particles, and experimental validation of CryoSiam particle identification by subtomogram averaging. **a**. Two-dimensional UMAP projections of embeddings generated by the SimSiam network for predicted particle instances in *M. pneumoniae* and *C. reinhardtii*. Each point represents one instance embedding, colored according to the log_10_ of its volume (number of voxels) in the predicted mask. **b**. The set of 14 macromolecular complexes used for the particle identification experiment. For each structure, the simulated density map is shown with the corresponding PDB ID indicated. Density maps were generated in ChimeraX from the atomic coordinates at 10 Å resolution. **c**. Initial *M. pneumoniae* subtomogram averaging results for labels with a sufficient number of predicted particles (>200 picks). Shown are ab initio models reconstructed in RELION (grey) and the refined subtomogram averages from SUSAN (blue) obtained using the ab initio models as references. **d**. Equivalent subtomogram averaging results for *C. reinhardtii* using the same reconstruction and refinement procedure as in panel c for two labels (label 1 and label 5).

