## Supplementary Material for "CryoSiam: self-supervised representation learning for automated analysis of cryo-electron tomograms"

### Supplementary Note 1 - CryoETSim dataset composition

**Supplementary Table 1:** Categories of macromolecules used in the individual samples in the CryoETSim dataset

| Sample | Simulated categories |
|---|---|
| general | macromolecules<br>cytoskeleton<br>membrane proteins<br>distractors<br>(smaller cytosolic complexes) |
| minimal bacterium | DNA filaments<br>transcription/translation complexes<br>membrane proteins<br>distractors |
| chromatin | DNA filaments<br>nucleosomes |

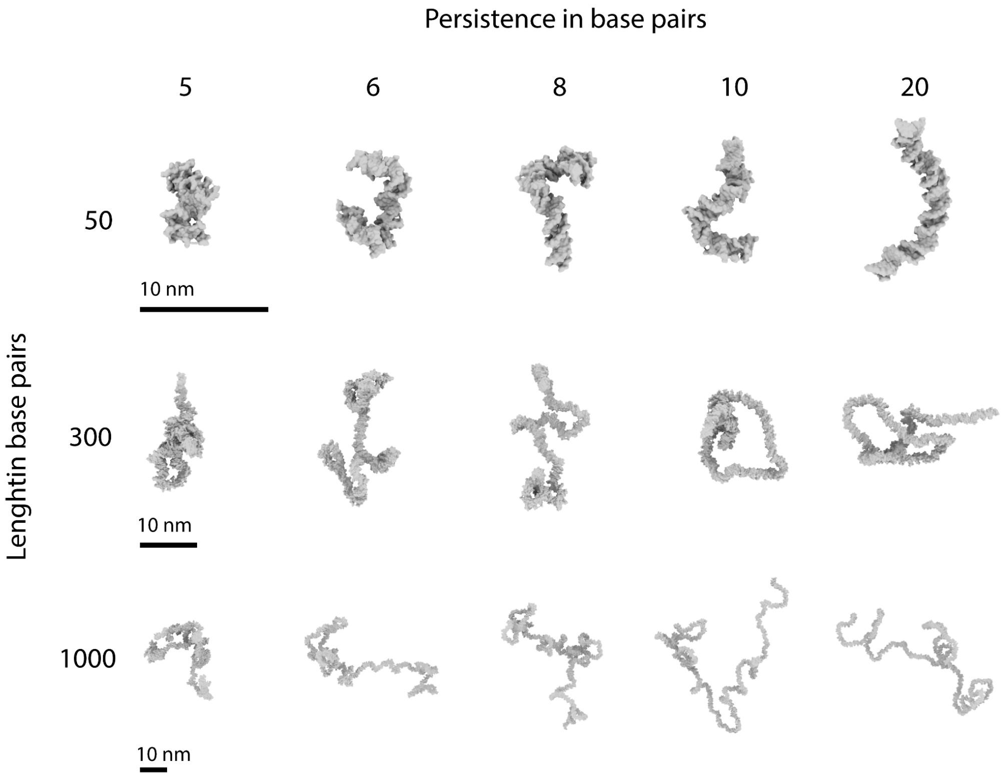

**Supplementary Figure 1: Simulation of short DNA filaments using the wormlike chain (WLC) model.** Example DNA filament fragments are shown for lengths of 50, 300, and 1000 base pairs (rows) and persistence lengths of 5-20 base pairs (columns). Increasing filament length produces more extended and flexible conformations, while higher persistence values result in smoother and less curved trajectories. Each filament was generated using the PolymerCpp software[1] and converted into the PDB file format for integration into the cryo-ET data simulation.

---

[1] Stefko, M., Douglass, K. & Manley, S. PolymerCpp. (Zenodo, 2020). doi:10.5281/zenodo.3928659.

**Supplementary Table 2:** Macromolecular complexes included in the simulated tomograms of the CryoETSim dataset

| Selection type | Particle name | Size (kDa) from PDB | PDB description |
|---|---|---|---|
| macromolecules | 7ELY | 2.90 | peptide |
| macromolecules | 7BLG | 18.14 | sugar binding protein |
| macromolecules | 3QM1 | 31.31 | serine cinnamoyl esterase |
| macromolecules | 1S3X | 42.75 | human Hsp70 ATPase |
| macromolecules | 6IOD | 55.33 | UdgX in complex with single-stranded DNA |
| macromolecules | 7S7K | 59.67 | EphB2 extracellular domain |
| macromolecules | 4WRM | 74.48 | human CSF-1:CSF-1R complex |
| macromolecules | 3H84 | 79.04 | GET3 chaperone |
| macromolecules | 3GL1 | 84.61 | ATPase domain of Ssb1 chaperone |
| macromolecules | 4UIC | 86.71 | S-layer protein rSbsC, sugar binding protein |
| macromolecules | 7E6G | 122.34 | diguanylate cyclase SiaD in complex with its activator SiaC |
| macromolecules | 6JYO | 135.63 | GII.13/21 noroviruses recognize glycans with a terminal beta-galactose via an unconventional glycan binding site |
| macromolecules | 7VTG | 136.04 | Pseudouridine kinase (PUKI) S30A mutant from Escherichia coli strain B |
| macromolecules | 7B5S | 166.62 | Ubiquitin ligation to F-box protein substrates by SCF-RBR E3-E3 super-assembly: CUL1-RBX1-ARIH1 Ariadne. Transition State 1 |
| macromolecules | 6lfm_gprotienpdb | 171.96 | |
| macromolecules | 7SHK | 172.48 | Xenopus laevis CRL2Lrr1 |

| | | | |
|---|---|---|---|
| macromolecules | 5LJO | 180.07 | E. coli BAM complex (BamABCDE), membrane protein |
| macromolecules | 2CG9 | 188.73 | Hsp90-Sba1 closed chaperone complex |
| macromolecules | 7SGM | 193.89 | Fab variant containing a fluorescent noncanonical amino acid with blocked excited state proton transfer and in complex with its antigen |
| macromolecules | 6n52_GluMpdb | 196.24 | |
| macromolecules | 5A20 | 197.50 | bacteriophage SPP1 head-to-tail interface filled with DNA and tape measure protein |
| macromolecules | 6ZIU | 197.83 | bovine ATP synthase stator domain, state 3 hydrolase |
| macromolecules | 6VGR | 203.29 | Human Dipeptidase 3 in Complex with Fab of SC-003 hydrolase |
| macromolecules | 2R9R | 206.88 | membrane/transport protein |
| macromolecules | 6LX3 | 208.31 | human secretory immunoglobulin A |
| macromolecules | 7WBT | 209.00 | bovine NLRP9 |
| macromolecules | 5CSA | 209.23 | BT-BCCP-AC1-AC5 of yeast acetyl-CoA carboxylase, ligase |
| macromolecules | 1UL1 | 214.07 | FEN1-PCNA complex, hydrolase/dna-binding protein |
| macromolecules | 3ULV | 226.08 | human TLR3ecd with three Fabs |
| macromolecules | 5JH9 | 230.78 | prApe1 hydrolase |
| macromolecules | 3D2F | 236.10 | Crystal structure of a complex of Sse1p and Hsp70, chaperone |
| macromolecules | 7NIU | 238.79 | Nanodisc reconstituted human ABCB4 in complex with 4B1-Fab and QA2-Fab |
| macromolecules | 1U6G | 238.82 | Cand1-Cul1-Roc1 Complex, ligase |

| | | | |
|---|---|---|---|
| macromolecules | 7BLR | 241.79 | Vps5 (SNX-BAR) complex |
| macromolecules | 2RHS | 245.45 | PheRS, ligase |
| macromolecules | 2WW2 | 252.90 | Family GH92 Inverting Mannosidase BT2199 from Bacteroides thetaiotaomicron VPI-5482 |
| macromolecules | 1N9G | 255.99 | thioester reductase, hydrolase |
| macromolecules | 6his_5HTpdb | 268.34 | |
| macromolecules | 3CF3 | 270.87 | Structure of P97/vcp in complex with ADP |
| macromolecules | 7KFE | 277.45 | Bundibugyo virus GP (mucin deleted) bound to antibody Fab BDBV-329 |
| macromolecules | 6ZQJ | 279.60 | trimeric prME spike of Spondweni virus |
| macromolecules | 1BXN | 279.81 | rubisco lyase |
| macromolecules | 6KRK | 288.88 | Peroxiredoxin from Aeropyrum pernix K1 |
| macromolecules | 6LMT | 295.89 | Cryo-EM structure of the killifish CALHM1 |
| macromolecules | 1QVR | 296.68 | ClpB, chaperone |
| macromolecules | 5H0S | 297.39 | VP1A and VP1B |
| macromolecules | 7JSN | 304.61 | Visual Signaling Complex between Transducin and Phosphodiesterase 6 |
| macromolecules | 7K5X | 311.14 | chromatosome containing human linker histone H1.0 |
| macromolecules | 6WZT | 323.80 | influenza hemagglutinin A/Victoria/361/2011 in complex with cyno antibody 3B10 |
| macromolecules | 3MKQ | 333.41 | yeast alpha/betaprime-COP subcomplex of the COPI vesicular coat |
| macromolecules | 6LXK | 352.70 | Z2B3 D102R Fab in complex with influenza virus neuraminidase from A/Serbia/NS-601/2014 |

| | | | |
|---|---|---|---|
| macromolecules | 6Z80 | 360.87 | stimulatory human GTP cyclohydrolase I - GFRP complex |
| macromolecules | 6CES | 370.31 | GATOR1-RAG |
| macromolecules | 2XNX | 371.16 | BC1 fragment of streptococcal M1 protein in complex with human fibrinogen |
| macromolecules | 6KSP | 382.87 | Rat GluD1 receptor(splayed conformation) in complex with 7-CKA and Calcium ions |
| macromolecules | 1SS8 | 386.04 | GroEL |
| macromolecules | 6VN1 | 392.96 | glycoprotein B-Neutralizing Antibody Complex |
| macromolecules | 6PIF | 421.50 | V. cholerae TniQ-Cascade complex, RNA bindng protein |
| macromolecules | 2DFS | 453.49 | Myosin-V, transport protein |
| macromolecules | 6CNJ | 470.11 | 2alpha3beta stiochiometry of the human Alpha4Beta2 nicotinic receptor |
| macromolecules | 6X5Z | 485.51 | Bovine Cardiac Myosin in Complex with Chicken Skeletal Actin and Human Cardiac Tropomyosin in the Rigor State |
| macromolecules | 6AHU | 492.83 | Ribonuclease P with mature tRNA |
| macromolecules | 7KJ2 | 495.88 | SARS-CoV-2 Spike Glycoprotein with one ACE2 Bound |
| macromolecules | 6U8Q | 511.64 | HIV-1 cleaved synaptic complex (CSC) intasome |
| macromolecules | 6KLH | 513.21 | Dimeric structure of Machupo virus polymerase bound to vRNA promoter |
| macromolecules | 3LUE | 541.23 | alpha-actinin CH1 bound to F-actin |
| macromolecules | 2VZ9 | 548.04 | Mammalian Fatty Acid Synthase in complex with NADP |

| | | | |
|---|---|---|---|
| macromolecules | 7B7U | 553.88 | mammalian RNA polymerase II in complex with human RPAP2 |
| macromolecules | 5O32 | 569.31 | complement complex, immune system |
| macromolecules | 6M04 | 595.87 | human homo-hexameric LRRC8D channel |
| macromolecules | 6TGC | 602.54 | ternary DOCK2-ELMO1-RAC1 complex |
| macromolecules | 7SFW | 606.58 | Venezuelan Equine Encephalitis virus (VEEV) TC-83 strain VLP in complex with Fab hVEEV-63 |
| macromolecules | 6BQ1 | 607.18 | Human PI4KIIIa lipid kinase complex |
| macromolecules | 6SCJ | 617.95 | human thyroglobulin |
| macromolecules | 7NYZ | 618.62 | MukBEF-MatP-DNA monomer |
| macromolecules | 7R04 | 635.36 | Neurofibromin in open conformation |
| macromolecules | 7NHS | 644.57 | Wzc K540M C8 |
| macromolecules | 7E8H | 653.34 | Kv4.2-DPP6S-KChIP1 complex |
| macromolecules | 6TAV | 666.20 | endopeptidase-induced alpha2-macroglobulin |
| macromolecules | 6XF8 | 689.88 | DLP 5 fold |
| macromolecules | 7AMV | 698.63 | poxvirus transcription pre-initiation complex, vRNAP |
| macromolecules | 7DD9 | 715.19 | Ams1 and Nbr1 complex, hydrolase, S. pombe |
| macromolecules | 6Z3A | 721.57 | Mec1-Ddc2 (wild-type) in complex with AMP-PNP |
| macromolecules | 6TA5 | 733.11 | OprM-MexA complex from the MexAB-OprM Pseudomonas aeruginosa whole assembly reconstituted in nanodiscs |

| | | | |
|---|---|---|---|
| macromolecules | 7Q21 | 735.45 | III2-IV2 respiratory supercomplex from Corynebacterium glutamicum |
| macromolecules | 6LXV | 751.32 | phosphoketolase from Bifidobacterium longum |
| macromolecules | 7KDV | 759.89 | Murine core lysosomal multienzyme complex (LMC) composed of acid beta-galactosidase (GLB1) and protective protein cathepsin A (PPCA, CTSA). hydrolase |
| macromolecules | 7LSY | 762.36 | NHEJ Short-range synaptic complex |
| macromolecules | 5FMG | 764.27 | Proteasome |
| macromolecules | 7EGQ | 801.37 | Coupling of N7-methyltransferase and 3'-5' exoribonuclease with polymerase reveals mechanisms for capping and proofreading |
| macromolecules | 1G3I | 826.23 | protease-chaperone complex |
| macromolecules | 6W6M | 874.43 | V. cholerae Type IV competence pilus secretin PilQ |
| macromolecules | 6TPS | 880.41 | early intermediate RNA Polymerase I Pre-initiation complex - eiPIC |
| macromolecules | 7O01 | 918.44 | Dimeric Photosystem I of a temperature sensitive mutant Chlamydomonas reinhardtii |
| macromolecules | 7ETM | 943.62 | C6 portal vertex in the enveloped virion capsid |
| macromolecules | 6MRC | 944.43 | ADP-bound human mitochondrial Hsp60-Hsp10 football complex, chaperone |
| macromolecules | 6VZ8 | 954.61 | arabidopsis thaliana acetohydroxyacid synthase complex with valine bound |
| macromolecules | 6KS8 | 968.94 | TRiC |

| | | | |
|---|---|---|---|
| macromolecules | 6EMK | 1041.14 | Saccharomyces cerevisiae Target of Rapamycin Complex 2 |
| macromolecules | 4XK8 | 1046.38 | photosystem I-LHCI super-complex |
| macromolecules | 7EEP | 1048.62 | Cyanophage Pam1 portal-adaptor complex |
| macromolecules | 7BKC | 1070.10 | Formate dehydrogenase - heterodisulfide reductase - formylmethanofuran dehydrogenase complex |
| macromolecules | 6GY6 | 1117.80 | XaxAB pore complex from Xenorhabdus nematophila |
| macromolecules | 6Z6O | 1144.83 | HDAC-TC |
| macromolecules | 7T3U | 1200.59 | IP3 ATP, and Ca2+ bound type 3 IP3 receptor in the inactive state |
| macromolecules | 7MEI | 1213.88 | Composite structure of EC+EC, transcription |
| macromolecules | 7QJ0 | 1224.45 | recombinant human gamma-Tubulin Ring Complex 6-spoked assembly intermediate |
| macromolecules | 5G04 | 1250.92 | human APC-Cdc20-Hsl1 complex |
| macromolecules | 7EGE | 1281.07 | TFIID in canonical conformation |
| macromolecules | 7WOO | 1286.20 | inner ring protomer of the Saccharomyces cerevisiae nuclear pore complex |
| macromolecules | 7SN7 | 1291.21 | enteropathogenic E. coli O127:H6 flagellar filament |
| macromolecules | 6GYM | 1297.09 | Structure of a yeast closed complex with distorted DNA - RNA polymerase II (Pol II) pre-initiation complex (PIC) |
| macromolecules | 4CR2 | 1309.28 | 26S proteasome |

| | | | |
|---|---|---|---|
| macromolecules | 7EGD | 1379.49 | SCP promoter-bound TFIID-TFIIA in initial TBP-loading state |
| macromolecules | 6YT5 | 1410.42 | T7 bacteriophage DNA translocation gp15-gp16 core complex intermediate assembly |
| macromolecules | 6DUZ | 1746.11 | periplasmic domains of PrgH and PrgK from the assembled Salmonella type III secretion injectisome needle complex, membrane protein |
| macromolecules | 6UP6 | 1797.10 | Endophilin B1 helical scaffold |
| macromolecules | 5OOL | 1797.38 | human mitochondrial ribosome with unfolded interfacial rRNA |
| macromolecules | 6F8L | 1818.38 | Thermus thermophilus PilF ATPase |
| macromolecules | 7EY7 | 2040.07 | bacteriophage T7 tail complex |
| macromolecules | 6ID1 | 2147.52 | human intron lariat spliceosome after Prp43 loaded |
| macromolecules | 6IGC | 2365.70 | HPV58/33/52 chimeric L1 pentamer |
| macromolecules | 6X9Q | 2796.28 | transcription-translation complex B3 (TTC-B3), ribosome, rna polymerase |
| macromolecules | 5MRC | 3325.59 | yeast mitochondrial ribosome |
| macromolecules | 4UJD | 4044.60 | mammalian 80S ribosome |
| cytoskeleton | MT_6o2tx2 | / | |
| cytoskeleton | actin cofilactin | / | |
| cytoskeleton | actin long | / | |
| cytoskeleton | neuroMT | / | |
| membrane proteins | 5SOA | 88.15 | Pseudomonas Aeruginosa FabF-C164Q mutant protein in complex with Z1613492358 |
| membrane proteins | GPROTEIN | 146.08 | G protein |

| | | | |
|---|---|---|---|
| membrane proteins | 6LFM_GPROTEIN | 171.96 | GPCR |
| membrane proteins | 6N52_GLUM | 196.24 | Metabotropic Glutamate Receptor 5 Apo Form |
| membrane proteins | 4COF_GABAAR | 209.20 | GABA(A)R-beta3 homopentamer |
| membrane proteins | 6HIS_5HT | 268.34 | Mouse serotonin 5-HT3 receptor |
| membrane proteins | 6WHV_GLUN1 | 416.39 | GluN1b-GluN2B NMDA receptor in complex with SDZ 220-040 and L689,560 |
| distractors | 1EXR | 16.89 | CA+2 bound calmodulin |
| distractors | 1QTX | 19.10 | Calmodulin RS20 peptide complex |
| distractors | 2Q0U | 43.72 | Pectenotoxin-2 and Latrunculin B Bound to Actin |
| distractors | 12GS | 48.01 | Glutathione S-transferase complexed with S-nonyl-glutathione |
| distractors | 1SJJ | 199.75 | Chicken Gizzard Smooth Muscle alpha-Actinin |
| transcription/translation | 1MWS | 149.62 | nitrocefin acyl-Penicillin binding protein 2a from methicillin resistant Staphylococcus aureus strain |
| transcription/translation | 6N60 | 458.37 | RNA polymerase sigma70-holoenzyme bound to upstream fork promoter DNA |
| transcription/translation | 6CA0 | 491.43 | E. coli RNAP sigma70 open complex |
| transcription/translation | 7OOC | 810.14 | 30S subunit of ribosomes in chloramphenicol-treated cells, Mycoplasma |
| transcription/translation | 1I94 | 841.86 | small ribosomal subunit with tetracycline, edeine and IF3 |
| transcription/translation | 4yg2 | 920.17 | RNA polymerase sigma70 holoenzyme |

| | | | |
|---|---|---|---|
| transcription/translation | 6AWB | 1141.70 | 30S ribosomal subunit and RNA polymerase complex in non-rotated state |
| transcription/translation | 7PAT | 1411.99 | free 50S in untreated Mycoplasma pneumoniae cells |
| transcription/translation | 7P6Z | 2250.56 | 70S ribosome in untreated cells, mycoplasma |
| transcription/translation | 4V4R | 2277.81 | whole ribosomal complex |
| transcription/translation | 6ZTN | 2669.95 | E. coli 70S-RNAP expressome complex in NusG-coupled state |
| transcription/translation | 4V5D | 4516.21 | 70S ribosome in complex with mRNA, paromomycin, acylated A- and P-site tRNAs, and E-site tRNA |
| transcription/translation | 4V5C | 4519.39 | 70S ribosome in complex with mRNA, paromomycin, acylated A-site tRNA, deacylated P-site tRNA, and E-site tRNA |
| transcription/translation | 4V5G | 4668.79 | 70S ribosome bound to EF-Tu and tRNA |
| transcription/translation | 4V5F | 4772.30 | ribosome with elongation factor G trapped in the post-translocational state |
| DNA filaments | dna_len_50_persis_5 | / | |
| DNA filaments | dna_len_50_persis_6 | / | |
| DNA filaments | dna_len_50_persis_8 | / | |
| DNA filaments | dna_len_50_persis_10 | / | |
| DNA filaments | dna_len_50_persis_20 | / | |
| DNA filaments | dna_len_100_persis_5 | / | |
| DNA filaments | dna_len_100_persis_6 | / | |
| DNA filaments | dna_len_100_persis_8 | / | |
| DNA filaments | dna_len_100_persis_10 | / | |
| DNA filaments | dna_len_100_persis_20 | / | |
| DNA filaments | dna_len_150_persis_5 | / | |
| DNA filaments | dna_len_150_persis_6 | / | |

| | | |
|---|---|---|
| DNA filaments | dna_len_150_persis_8 | / |
| DNA filaments | dna_len_150_persis_10 | / |
| DNA filaments | dna_len_150_persis_20 | / |
| DNA filaments | dna_len_200_persis_5 | / |
| DNA filaments | dna_len_200_persis_6 | / |
| DNA filaments | dna_len_200_persis_8 | / |
| DNA filaments | dna_len_200_persis_10 | / |
| DNA filaments | dna_len_200_persis_20 | / |
| DNA filaments | dna_len_250_persis_5 | / |
| DNA filaments | dna_len_250_persis_6 | / |
| DNA filaments | dna_len_250_persis_8 | / |
| DNA filaments | dna_len_250_persis_10 | / |
| DNA filaments | dna_len_250_persis_20 | / |
| DNA filaments | dna_len_300_persis_5 | / |
| DNA filaments | dna_len_300_persis_6 | / |
| DNA filaments | dna_len_300_persis_8 | / |
| DNA filaments | dna_len_300_persis_10 | / |
| DNA filaments | dna_len_300_persis_20 | / |
| DNA filaments | dna_len_350_persis_5 | / |
| DNA filaments | dna_len_350_persis_6 | / |
| DNA filaments | dna_len_350_persis_8 | / |
| DNA filaments | dna_len_350_persis_10 | / |
| DNA filaments | dna_len_350_persis_20 | / |
| DNA filaments | dna_len_400_persis_5 | / |
| DNA filaments | dna_len_400_persis_6 | / |
| DNA filaments | dna_len_400_persis_8 | / |
| DNA filaments | dna_len_400_persis_10 | / |
| DNA filaments | dna_len_400_persis_20 | / |
| DNA filaments | dna_len_450_persis_5 | / |
| DNA filaments | dna_len_450_persis_6 | / |
| DNA filaments | dna_len_450_persis_8 | / |
| DNA filaments | dna_len_450_persis_10 | / |
| DNA filaments | dna_len_450_persis_20 | / |

| | | | |
|---|---|---|---|
| DNA filaments | dna_len_500_persis_5 | / | |
| DNA filaments | dna_len_500_persis_6 | / | |
| DNA filaments | dna_len_500_persis_8 | / | |
| DNA filaments | dna_len_500_persis_10 | / | |
| DNA filaments | dna_len_500_persis_20 | / | |
| DNA filaments | dna_len_1000_persis_5 | / | |
| DNA filaments | dna_len_1000_persis_6 | / | |
| DNA filaments | dna_len_1000_persis_8 | / | |
| DNA filaments | dna_len_1000_persis_10 | / | |
| DNA filaments | dna_len_1000_persis_20 | / | |
| nucleosomes | 5F99 | 199.29 | MMTV-A Nucleosome Core Particle |
| nucleosomes | 7PEY | 219.68 | Nucleosome 3 of the 4x177 nucleosome array containing H1 |
| nucleosomes | 7PEW | 222.77 | Nucleosome 1 of the 4x177 nucleosome array containing H1 |
| nucleosomes | 7PEX | 245.19 | Nucleosome 2 of the 4x177 nucleosome array containing H1 |
| nucleosomes | 5OXV | 411.66 | 4_601_157 tetranucleosome |
| nucleosomes | 1ZBB | 431.87 | 4_601_167 Tetranucleosome |
| nucleosomes | 6M3V | 439.12 | 355 bp di-nucleosome harboring cohesive DNA termini |
| nucleosomes | 6LA8 | 457.42 | 349 bp di-nucleosome harboring cohesive DNA termini assembled with linker histone H1.0 |
| nucleosomes | 6L4A | 635.98 | H3-H3-H3 tri-nucleosome with the 22 base-pair linker DNA |
| nucleosomes | 6L49 | 637.16 | H3-CA-H3 tri-nucleosome with the 22 base-pair linker DNA |
| nucleosomes | 7PEV | 661.93 | 4x177 nucleosome array containing H1 |

| | | | |
|---|---|---|---|
| nucleosomes | 7PEU | 685.97 | 4x177 nucleosome array containing H1 |
| nucleosomes | 7PF0 | 720.12 | Trinucleosome of the 4x177 nucleosome array containing H1 |
| nucleosomes | 7VA4 | 745.81 | Telomeric tetranucleosome in open state |
| nucleosomes | 7V9K | 750.15 | Telomeric tetranucleosome |
| nucleosomes | 7PFT | 772.82 | Trinucleosome of the 4x207 nucleosome array containing H1 |
| nucleosomes | 5OY7 | 827.37 | 4_601_157 tetranucleosome |
| nucleosomes | 7PFA | 872.74 | Trinucleosome of the 4x197 nucleosome array containing H1 |
| nucleosomes | 7PET | 933.73 | 4x177 nucleosome array containing H1 |

### Supplementary Note 2 - Training of CryoSiam on the CryoETSim simulated dataset

**Supplementary Table 3:** Train and test tomogram sets for downstream tasks of DenseSimSiam on the CryoETSim dataset

| Downstream task | Sample type | Train dataset | Test dataset |
|---|---|---|---|
| Denoising | general | sample_1-sample_200 | sample_201-sample_299 |
| Denoising | minimal_bacterium | sample_301-sample_340 | sample_341-sample_349 |
| Denoising | chromatin | sample_350-sample_390 | sample_391-sample_400 |
| Semantic segmentation | general | sample_1-sample_200 | sample_201-sample_299 |
| Semantic segmentation | minimal_bacterium | sample_301-sample_340 | sample_341-sample_349 |
| Semantic segmentation | chromatin | sample_350-sample_390 | sample_391-sample_400 |
| Instance segmentation | general | sample_1-sample_200 | sample_200-sample_300 |

**Supplementary Table 4:** Evaluation of DenseSimSiam on the downstream task of semantic segmentation

| Classes | Clean | Noisy | Denoised |
|---|---|---|---|
| background | 0.944 | 0.894 | 0.896 |
| membrane | 0.778 | 0.639 | 0.671 |
| microtubules | 0.874 | 0.754 | 0.793 |
| particles | 0.822 | 0.619 | 0.636 |
| actin | 0.788 | 0.280 | 0.342 |
| DNA filaments | 0.703 | 0.332 | 0.350 |
| all | 0.836 | 0.635 | 0.664 |

**Supplementary Table 5:** Evaluation of DenseSimSiam on the downstream task of instance segmentation with average prediction (AP) applying different Intersection over Union (IoU) thresholds (50, 55, 60, 65, 70, 75, 80).

| AP | Clean | Noisy | Denoised |
|---|---|---|---|
| AP-50 | 0.437 | 0.229 | 0.249 |
| AP-55 | 0.417 | 0.180 | 0.196 |
| AP-60 | 0.393 | 0.121 | 0.133 |
| AP-65 | 0.346 | 0.063 | 0.071 |
| AP-70 | 0.241 | 0.020 | 0.024 |
| AP-75 | 0.099 | 0.003 | 0.003 |
| AP-80 | 0.016 | 0.000 | 0.000 |

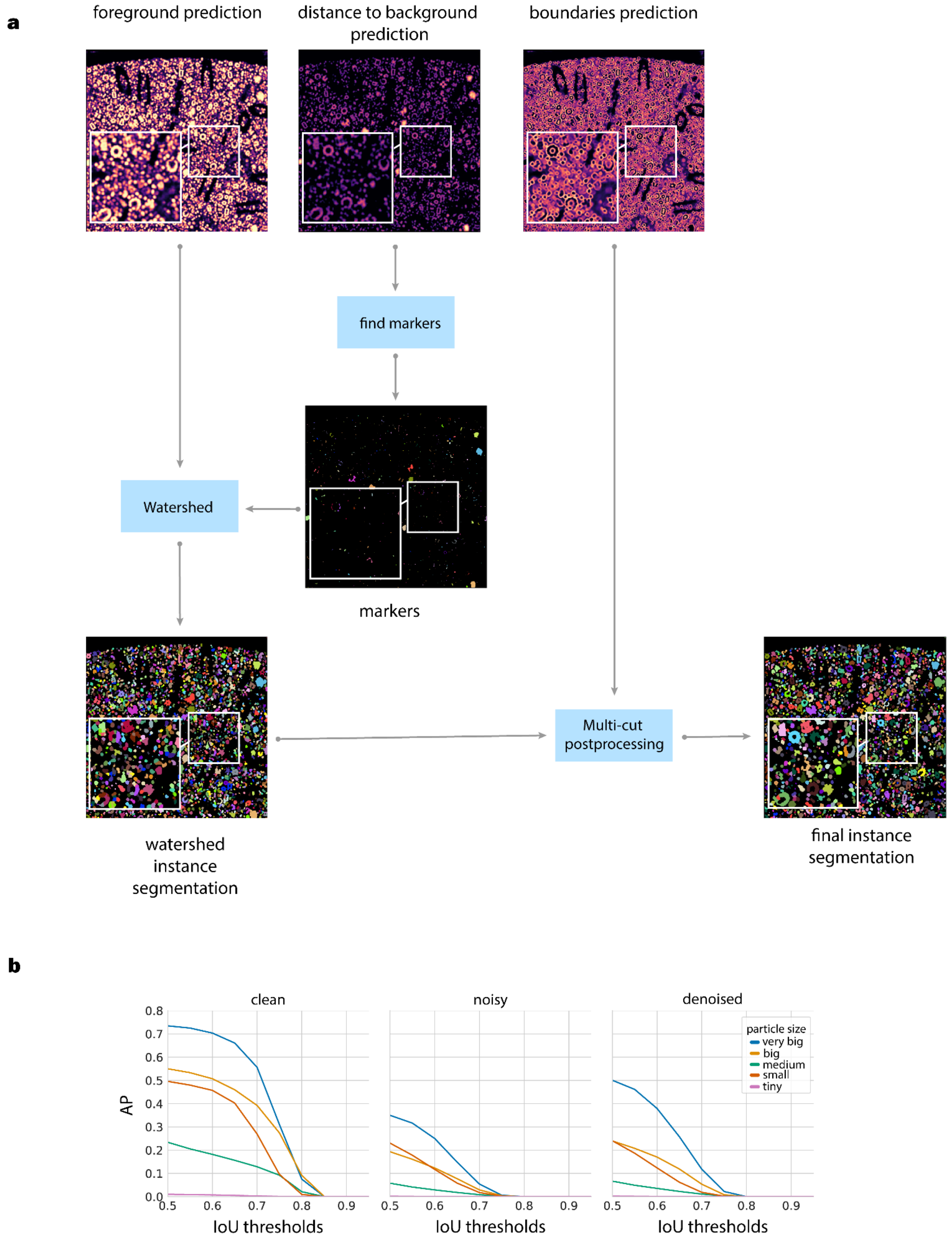

**Supplementary Figure 2: Overview of the instance segmentation workflow.**

**a.** Overview of the instance segmentation workflow. The network predicts three voxel-wise components: foreground regions (voxels belonging to macromolecular complexes/particles), distance-to-background maps (voxel-wise distances from the nearest background region), and instance boundaries (interfaces between adjacent particles). Particle markers are extracted from the distance maps via Gaussian filtering and thresholding. The watershed algorithm[2] combines the predicted foreground map and extracted markers to generate preliminary instance segmentation masks. These masks are further refined using multi-cut postprocessing[3,4], which incorporates the predicted boundary map to correct over-segmentation and improve particle delineation. Insets highlight densely crowded regions showing separation of adjacent instances.

**b.** Average precision (AP) curves across different intersection-over-union (IoU) thresholds for clean, noisy, and denoised tomograms. Each curve corresponds to a particle size category (very big, big, medium, small, tiny). Larger particles achieve higher AP scores across all conditions, while smaller particles show greater sensitivity to noise and contrast loss.

[2] Beucher, S. The watershed transformation applied to image segmentation. Scanning Microsc. 1992, 28 (1992).

[3] Kernighan, B. W. & Lin, S. An efficient heuristic procedure for partitioning graphs. Bell Syst. Tech. J. 49, 291–307 (1970).

[4] Wolf, S. et al. The Mutex Watershed and its Objective: Efficient, Parameter-Free Graph Partitioning. IEEE Trans. Pattern Anal. Mach. Intell. 43, 3724–3738 (2021).

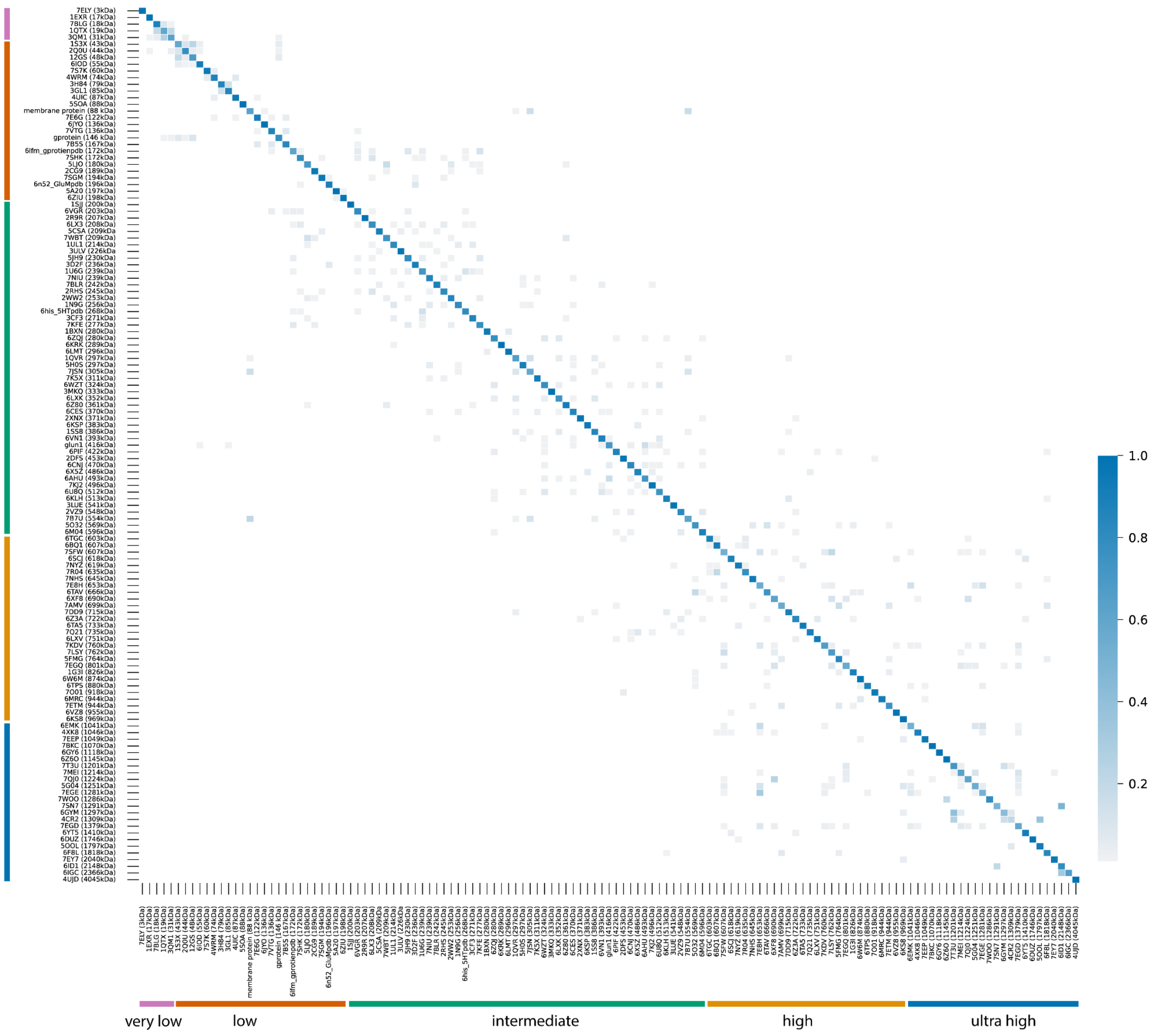

**Supplementary Figure 3: Confusion matrix from the kNN classifier with embeddings from SimSiam finetuned with contrastive learning on noisy simulated data.** Each entry shows the proportion of particles from a true class/particle identity (rows) that were predicted as a given class (columns). The matrix illustrates the consistency in classification between molecular identities based on their learned representations. Rows and columns are ordered by increasing molecular weight (kDa), and each row is normalized to span values from 0 to 1.

### Supplementary Note 3 - Ablation studies

**Supplementary Table 6:** Train and test tomogram sets used for the ablation studies. A total of 48 tomograms were included in the training set and 12 tomograms in the test set. For each ablation experiment, the DenseSimSiam model was first trained in a self-supervised manner, after which a simple three-layer CNN segmentation head was added to evaluate the effect of each configuration on semantic segmentation performance.

| Dataset type | Simulated data sample type | Selected tomograms |
|---|---|---|
| Train | general | sample_1-sample_32 |
| Train | minimal bacterium | sample_301-sample_308 |
| Train | chromatin | sample_351-sample_358 |
| Test | general | sample_33-sample_40 |
| Test | minimal bacterium | sample_309-sample_310 |
| Test | chromatin | sample_359-sample_360 |

**Supplementary Table 7:** Results of the ablation study evaluating the effect of different transformations on DenseSimSiam performance. The table reports semantic segmentation accuracy across structural classes (background, membranes, microtubules, particles, actin, and DNA filaments) under four transformation settings: no transformations, Gaussian low-pass filtering, combined low-pass and high-pass filtering, and the addition of Gaussian noise. The results demonstrate that combining multiple transformations (low-pass, high-pass, and Gaussian noise) yields the most robust and generalizable segmentation performance.

| Classes | None | Low-pass | Low-pass, High-pass | Low-pass, High-pass, Gaussian Noise |
|---|---|---|---|---|
| background | 0.835 | 0.614 | 0.900 | 0.873 |
| membrane | 0.000 | 0.000 | 0.000 | 0.255 |
| microtubules | 0.000 | 0.000 | 0.000 | 0.000 |
| particles | 0.525 | 0.562 | 0.599 | 0.604 |
| actin | 0.000 | 0.016 | 0.000 | 0.027 |
| dna filaments | 0.003 | 0.299 | 0.374 | 0.328 |
| all | 0.302 | 0.286 | 0.378 | 0.416 |

**Supplementary Table 8:** Results of the ablation study evaluating the effect of voxel masking during self-supervised training. The table reports Dice scores for different masking ratios (0%, 10%, 25%, 50%, 75%, and 90%). Although the model trained without masking (0%) achieved the highest overall scores, a 50% masking ratio provided the best balance between robustness and accuracy, particularly improving the segmentation of particles and DNA filaments.

| Classes | 0% | 10% | 25% | 50% | 75% | 90% |
|---|---|---|---|---|---|---|
| background | 0.860 | 0.856 | 0.832 | 0.870 | 0.531 | 0.147 |
| membrane | 0.236 | 0.000 | 0.203 | 0.000 | 0.161 | 0.017 |
| microtubules | 0.166 | 0.087 | 0.000 | 0.000 | 0.235 | 0.013 |
| particles | 0.589 | 0.532 | 0.542 | 0.615 | 0.484 | 0.406 |
| actin | 0.000 | 0.000 | 0.000 | 0.000 | 0.030 | 0.013 |
| dna filaments | 0.351 | 0.302 | 0.252 | 0.384 | 0.132 | 0.306 |
| all | 0.431 | 0.363 | 0.376 | 0.379 | 0.286 | 0.153 |

**Supplementary Table 9:** Results of the ablation study assessing the effect of voxel embedding dimensionality on segmentation performance. The table reports Dice scores for models trained with voxel embedding sizes of 8, 16, 32, and 64 dimensions. The 64-dimensional embeddings achieved the highest overall performance, providing a strong balance between representational richness and generalization across structural classes.

| Classes | 8-d | 16-d | 32-d | 64-d |
|---|---|---|---|---|
| background | 0.841 | 0.808 | 0.000 | 0.887 |
| membrane | 0.000 | 0.000 | 0.137 | 0.252 |
| microtubules | 0.000 | 0.240 | 0.010 | 0.210 |
| particles | 0.567 | 0.556 | 0.526 | 0.620 |
| actin | 0.000 | 0.033 | 0.000 | 0.056 |
| dna filaments | 0.254 | 0.202 | 0.301 | 0.411 |
| all | 0.349 | 0.367 | 0.175 | 0.466 |

**Supplementary Table 10:** Results of the ablation study evaluating the effect of global embedding dimensionality on segmentation performance. The table reports Dice scores for models trained with global embedding sizes of 128, 256, 512, and 1024 dimensions. The 256-dimensional embeddings achieved the best overall performance, offering an optimal balance between structural feature representation and model stability.

| Classes | 128-d | 256-d | 512-d | 1024-d |
|---|---|---|---|---|
| background | 0.000 | 0.904 | 0.859 | 0.888 |
| membrane | 0.184 | 0.000 | 0.000 | 0.000 |
| microtubules | 0.007 | 0.000 | 0.000 | 0.000 |
| particles | 0.603 | 0.674 | 0.594 | 0.635 |
| actin | 0.044 | 0.000 | 0.000 | 0.000 |
| dna filaments | 0.298 | 0.537 | 0.288 | 0.393 |
| all | 0.205 | 0.416 | 0.364 | 0.389 |

**Supplementary Table 11**: Results of the ablation study examining the effect of different embedding loss combinations on segmentation performance. The table reports Dice scores for models trained with voxel-level (Dense) loss only, and with the addition of global and intermediate-level losses (L2, L4, and L8), corresponding to pooling factors of 2, 4, and 8, respectively. Incorporating the L2-level loss alongside dense and global losses yielded the best overall performance, providing an effective balance between fine-scale detail and global contextual representation.

| Classes | Dense | Global | L2 | L4 | L8 |
|---|---|---|---|---|---|
| background | 0.000 | 0.701 | 0.891 | 0.879 | 0.000 |
| membrane | 0.020 | 0.270 | 0.000 | 0.000 | 0.000 |
| microtubules | 0.095 | 0.202 | 0.342 | 0.000 | 0.256 |
| particles | 0.422 | 0.613 | 0.616 | 0.619 | 0.413 |
| actin | 0.015 | 0.039 | 0.113 | 0.000 | 0.000 |
| dna filaments | 0.212 | 0.313 | 0.428 | 0.308 | 0.292 |
| all | 0.132 | 0.402 | 0.452 | 0.375 | 0.159 |

### Supplementary Note 4 - CryoSiam on real data

**Supplementary Table 12**: Data acquisition parameters for the experimental tomograms of M. pneumoniae and C. reinhardtii used in this study. The table summarizes the imaging conditions for datasets obtained from EMPIAR (EMPIAR-10499 and EMPIAR-11756) and the CryoET Data Portal (datasets DS-10003 and DS-10301), including microscope type, detector, energy filter, magnification, dose per tilt, number of tilts, tilt increment, and defocus range.

| Parameter | *M. pneumoniae* | *C. reinhardtii* |
|---|---|---|
| Microscope | Titan Krios G3 | Titan Krios G4 |
| Detector | Gatan K2 Summit | Thermo Fisher Falcon 4i |
| Energy filter | Quantum post-column energy filter | Selectris X energy filter |
| Magnification | 81000x | 64000x |
| Pixel size [Å/pixel] | 1.7005 | 1.82/1.90 |
| Dose per tilt [$e^{-}$/Å$^{-2}$] | 3.24 | 1.64 |
| Number of Tilts | 41 | 40 |
| Tilt increment | 3° | 3° |
| Defocus range | 1.5-3.5 µm | 1.5-3.5 µm |

**Supplementary Table 13**: Summary of subtomogram averaging (STA) results from CryoSiam particle identification predictions. The table reports quantitative results from STA performed on predicted particle coordinates generated by CryoSiam. Columns list the total number of predicted instances per label, the number of particles contributing to the best ab initio class obtained in RELION, and final refinement results from the second STA stage, including the number of tomograms contributing particles, the total number of particles used, and the final resolution achieved after refinement.

| Organism | Particle | CryoSiam picks | Best class in ab initio | Final results from the second stage of STA | | |
|---|---|---|---|---|---|---|
| | | | | Number of tomograms | Final number of particles | Final resolution[5] |
| *M. pneumoniae* | label 1 | 11,023 | 5,190 | / | / | / |
| *M. pneumoniae* | label 4 | 6,642 | 3,165 | / | / | / |
| *M. pneumoniae* | label 5 | 37,218 | 16,781 | 65 | 17,681 | **13.6 Å** |
| *M. pneumoniae* | Label 5 (50S ribosomal subunit class) | / | / | 65 | 3,060 | 18.1 Å |
| *M. pneumoniae* | label 7 | 2,151 | 1,982 | / | / | / |
| *M. pneumoniae* | label 8 | 3,991 | 1,693 | / | / | / |
| *M. pneumoniae* | label 9 | 2,708 | 1,299 | 59 | 193 | 21.8 Å |
| *M. pneumoniae* | label 10 | 1,161 | 679 | / | / | / |
| *M. pneumoniae* | label 11 | 2,756 | 877 | / | / | / |
| *M. pneumoniae* | label 13 | 4,790 | 1,948 | 61 | 234 | 27.2 Å |
| *M. pneumoniae* | label 14 | 243 | 217 | / | / | / |
| *C. reinhardtii* | label 1 | 5,063 | 1,524 | / | / | / |
| *C. reinhardtii* | label 5 | 11,858 | 4,111 | 13 | 2,680 | **15.7 Å** |

[5] Bold value indicates that the resolution reached the Nyquist limit of extracted subtomograms

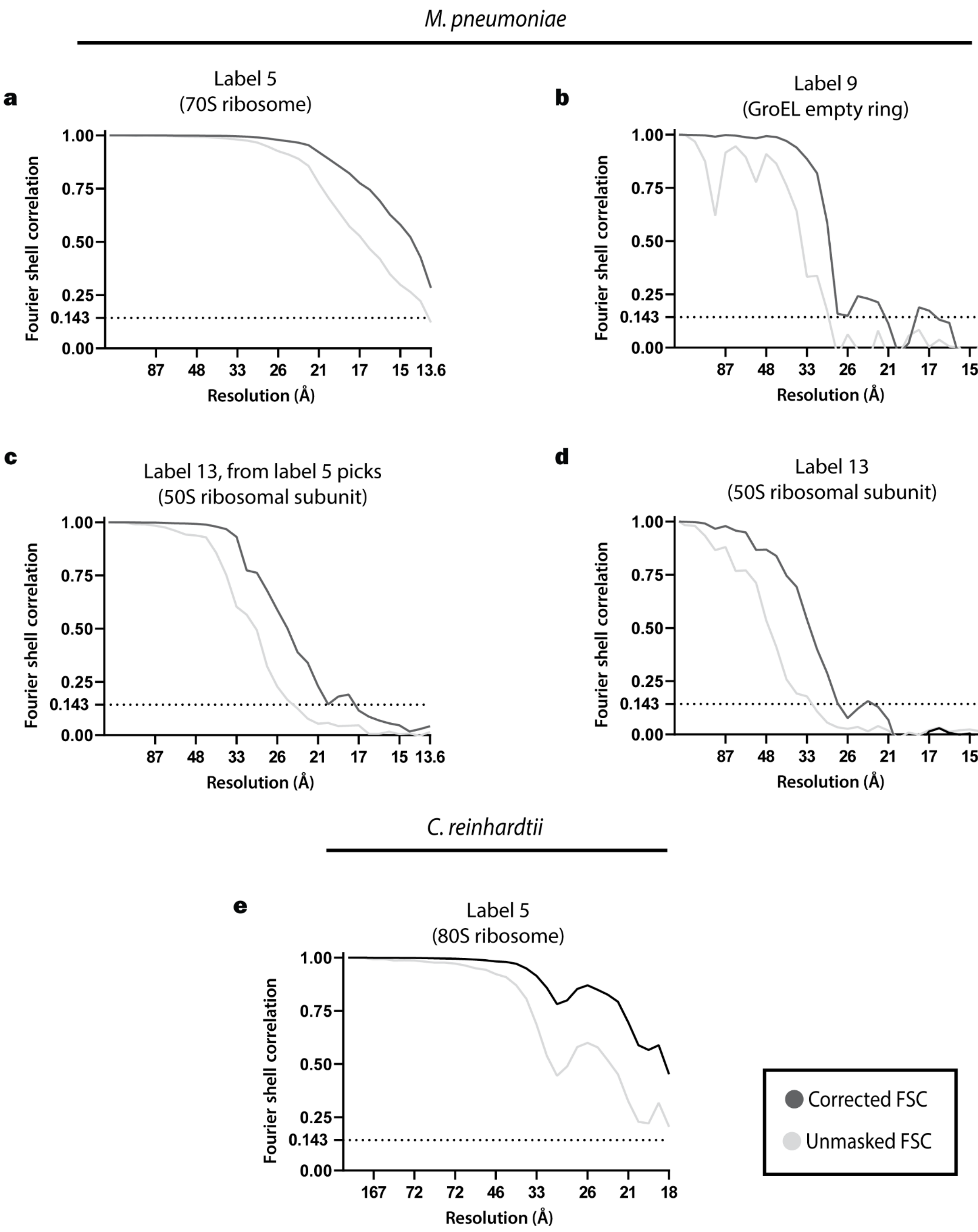

**Supplementary Figure 4: Fourier shell correlation (FSC) plots for subtomogram averaging results.** The FSC of the subtomogram average of the 70S ribosome (**a; label 5**), GroEL empty ring (**b; label 9**), 50S ribosomal subunit coming from particle picks from label 5 (**c; label 5**), and 50S ribosomal subunit (**d; label 13**) from *M. pneumoniae* (EMPIAR-10499) and the 80S ribosome (**e; label 5**) from *C. reinhardtii* (EMPIAR-11756). The corrected FSC is shown in dark grey, and the unmasked FSC is shown in light grey.
